# Predictions of immunogenicity reveal potent SARS-CoV-2 CD8+ T-cell epitopes

**DOI:** 10.1101/2022.05.23.492800

**Authors:** David Gfeller, Julien Schmidt, Giancarlo Croce, Philippe Guillaume, Sara Bobisse, Raphael Genolet, Lise Queiroz, Julien Cesbron, Julien Racle, Alexandre Harari

## Abstract

The recognition of pathogen or cancer-specific epitopes by CD8^+^ T cells is crucial for the clearance of infections and the response to cancer immunotherapy. This process requires epitopes to be presented on class I Human Leukocyte Antigen (HLA-I) molecules and recognized by the T-Cell Receptor (TCR). Machine learning models capturing these two aspects of immune recognition are key to improve epitope predictions. Here we assembled a high-quality dataset of naturally presented HLA-I ligands and experimentally verified neo-epitopes. We then integrated these data with new algorithmic developments to improve predictions of both antigen presentation and TCR recognition. Applying our tool to SARS-CoV-2 proteins enabled us to uncover several epitopes. TCR sequencing identified a monoclonal response in effector/memory CD8^+^ T cells against one of these epitopes and cross-reactivity with the homologous SARS-CoV-1 peptide.

## Introduction

CD8^+^ T cells have the ability to eliminate infected or malignant cells and play a key role in infectious diseases and cancer immunotherapy. CD8^+^ T-cell recognition is initiated by the binding of the T-cell Receptor (TCR) to peptides, referred to as epitopes, displayed on class I human leukocyte antigens (HLA-I) molecules. Detailed knowledge of class I epitopes in cancer and infectious diseases has several translational and clinical applications. Such epitopes can be used to design vaccines that target the most relevant epitopes, including neo-epitopes (i.e., peptides derived from non-synonymous genetic alterations) in cancer (Ott et al., 2017; Sahin et al., 2017, 2020). These epitopes can also be used to select TCRs, study them and reinfuse these TCRs into patients as part of adoptive T-cell therapy (Rosenberg and Restifo, 2015). Unfortunately, identifying epitopes in cancer or infectious diseases is challenging because of the very high number of possible candidates and the diversity of HLA-I alleles. For instance, for each non-synonymous point mutation in a tumor, up to 38 8- to 11-mer peptides containing the mutated residue may be immunogenic. Similarly, the number of potential class I epitopes of a given length in a pathogen is roughly equal to the length of the proteome of this pathogen. Major improvements have been done for experimentally screening potential epitope candidates, either with peptide pools (Tarke et al., 2021) or tandem mini-genes (Parkhurst et al., 2017; Tran et al., 2015). Nevertheless, the most common approach to identify new epitopes is to preselect them based on HLA-I ligand predictors.

HLA-I molecules are encoded by three genes (HLA-A, -B and -C). These genes are highly polymorphic in human and different alleles are characterized by specific binding motifs and specific peptide length distributions in their ligands (Gfeller and Bassani-Sternberg, 2018). Binding motifs mainly reflect amino acids favorable for binding to HLA-I molecules at specific positions of the ligands. Peptide length distributions (typically from 8- to 14-mers with a preference for 9-mers for most alleles) capture both the binding preferences of HLA-I molecules as well as the skewed length distribution of peptides available in the endoplasmic reticulum for loading onto HLA-I molecules (Trolle et al., 2016).

The specificity of HLA-I binding motifs and peptide length distributions greatly constrains the repertoire of potential epitopes. As such, computational tools that accurately capture these two features of antigen presentation have been developed to narrow down the list of potential epitopes to be experimentally tested. Historically, predictors HLA-I ligands were mainly trained on peptides tested experimentally in binding assays (Peters et al., 2020), with the caveat that many of these peptides had been pre-selected based on previous versions of the predictors. More recently, naturally presented HLA-I ligands identified by mass spectrometry (MS) based HLA-I peptidomics provided a rich source of information about the rules of antigen presentation and the specificity of HLA-I molecules (Abelin et al., 2017; Alvarez et al., 2019; Bassani-Sternberg and Gfeller, 2016; Bassani-Sternberg et al., 2017; Di Marco et al., 2017; Sarkizova et al., 2019). The number of HLA-I ligands identified by this technology, both in mono-allelic and poly-allelic samples, surpasses the one from binding assays and HLA-I peptidomics data are now included in the training of most HLA-I ligand predictors (Gfeller et al., 2018; O’Donnell et al., 2020; Pyke et al., 2021; Reynisson et al., 2020; Sarkizova et al., 2019; Shao et al., 2020). For poly-allelic samples, motif deconvolution has been used to identify HLA-I binding motifs and determine allelic restriction of HLA-I ligands without relying on HLA-I ligand predictors (Alvarez et al., 2019; Andreatta et al., 2017; Bassani-Sternberg and Gfeller, 2016).

Over the years, several attempts have been made to integrate additional features in epitope predictions linked to antigen presentation and TCR recognition. For instance, gene expression and protein abundance were shown to improve HLA-I ligand and class I epitope predictions (Abelin et al., 2017; Koşaloğlu-Yalçın et al., 2022; Sarkizova et al., 2019). Predictions of cleavage or antigen transport properties were explored (Stranzl et al., 2010). The concept of antigen presentation hotspot, as determined by the analysis of HLA-I peptidomics data, was also shown to improve predictions (Müller et al., 2017). Some studies further attempted to integrate TCR recognition propensities in epitope predictions, for instance by investigating the role of dissimilarity-to-self or foreignness (Balachandran et al., 2017; Duan et al., 2014; Łuksza et al., 2017; Wells et al., 2020). We and others observed that specific amino acids found in epitope residues more likely to interact with the TCR increase the propensity for TCR recognition (Calis et al., 2013; Chowell et al., 2015; Schmidt et al., 2021).

In this work, we compiled and curated a large dataset of HLA-I ligands and neoepitopes, and trained predictors of antigen presentation and TCR recognition. Applying these tools to SARS-CoV-2 proteins enabled us to predict and validate several epitopes, which we characterize in terms of TCR functional avidity, clonality and crossreactivity.

## Results

### Integration and curation of HLA-I peptidomics data reveal binding motifs and peptide length distributions for more than hundred alleles

To improve predictions of class I antigen presentation, we manually compiled recent studies of naturally presented HLA-I ligands profiled by mass spectrometry. Our dataset covers 24 studies for a total of 244 samples (see Dataset S1). All data were retrieved from the original publications and were not filtered by any HLA-I ligand predictor, ensuring that our dataset is not biased by such filtering. All samples were processed with the motif deconvolution tool MixMHCp and shared motifs across samples containing the same HLA-I allele were annotated to this allele, following our previously established approach (see example in Figure 1A) (Bassani-Sternberg and Gfeller, 2016; Bassani-Sternberg et al., 2017). All motifs in all samples were manually verified, and samples or alleles for which motif deconvolution results were ambiguous were not considered. This enabled us to derive reliable binding motifs and peptide length distributions for 119 HLA-I alleles, supported by a total of 384,070 peptides (Dataset S2, see examples in Figure 1B). Motifs for HLA-I alleles identified in mono-allelic and poly-allelic samples were highly similar (Figure S1). Peptide length distributions for alleles in mono-allelic samples displayed a slightly lower fraction of 9-mers and a slightly higher fraction of peptides of other lengths compared to those observed poly-allelic samples (Figure 1C).

**Figure 1:**
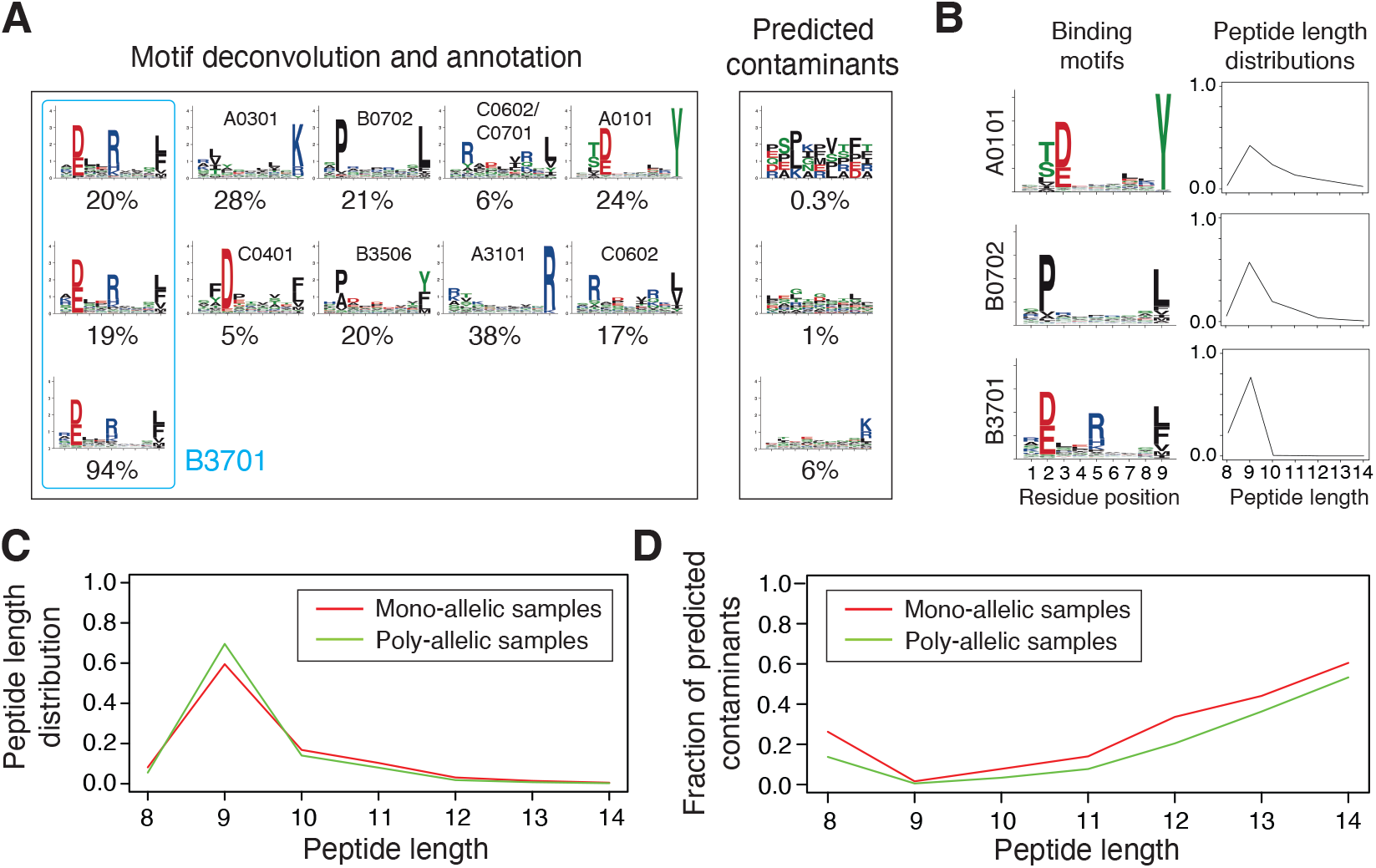
Integration and curation of HLA-I peptidomics data reveal binding motifs and peptide length distributions for more than hundred alleles. (A) Motif deconvolution includes identification of motifs and predicted contaminants with MixMHCp, as well as motif annotation by identifying shared motifs across samples sharing the same allele. The example shows the deconvolved motifs in two poly-allelic samples that share the HLA-B*37:01 allele (‘donor1’ and ‘HCC1143’ in Dataset S1), as well as the mono-allelic HLA-B*37:01 sample. (B) Examples of binding motifs and peptide length distributions obtained by motif deconvolution and used to train MixMHCpred2.2. (C) Peptide length distributions for alleles observed in both mono-allelic and poly-allelic HLA-I peptidomics data. Each curve represents the average peptide length distribution across these alleles. (D) Fraction of predicted contaminants across different lengths (average over all samples).

In addition to identifying binding motifs and peptide length distributions for the different alleles expressed in a sample, motif deconvolution is useful to remove predicted contaminants that can arise from wrongly identified spectra or experimental contaminations (Andreatta et al., 2017; Gfeller et al., 2018). As previously reported (Gfeller et al., 2018), the fraction of predicted contaminants is especially large in 8-mers and 11- to 14-mers both in mono-allelic and poly-allelic samples, representing for instance more than 50% of 14-mers (Figure 1D, see specific examples in Figure S2A). Predicted contaminants include peptides with trypsin-like motifs or from other HLA-I alleles (see examples in Figure 1A and Figure S2B). Motif deconvolution is also powerful to pinpoint putatively erroneous HLA-I typing (Figure S2C). These observations demonstrate the importance of performing careful quality-control before using HLA-I peptidomics data to train predictors (Fritsche et al., 2021).

### Models of HLA-I binding specificities and peptide length distributions improve predictions of naturally presented HLA-I ligands

To improve predictions of class I antigen presentation, we integrated these data into the training of our HLA-I ligand predictor MixMHCpred and further refined the modelling of peptide length distributions (see Methods). As with most HLA-I ligand predictors, the final score of a peptide is expressed as a %rank, which represents how the predicted binding of a peptide compares to the one of random peptides from the human proteome (see Methods). To benchmark the new version of MixMHCpred (v2.2) we used two external datasets. The first one consists of 10 HLA-I peptidomics datasets from meningioma samples (Gfeller et al., 2018). The second one consists of 11 recently published HLA-I peptidomics datasets (Pyke et al., 2021). These datasets were not included in the training of any predictor considered in this work. 4-fold excess of randomly selected peptides from the human proteome were used as negatives to compute Receiver Operating Curves (ROC) and Positive Predictive values (PPV) (see Methods). Both the Area Under the ROC Curve (AUC) and the PPV were higher for MixMHCpred2.2, compared to NetMHCpan4.1 (Reynisson et al., 2020), MHCflurry2.0 (O’Donnell et al., 2020), HLAthena (Sarkizova et al., 2019) and MixMHCpre2.0.2 (Gfeller et al., 2018) (Figure 2A-B and Figure S3A-D).

**Figure 2:**
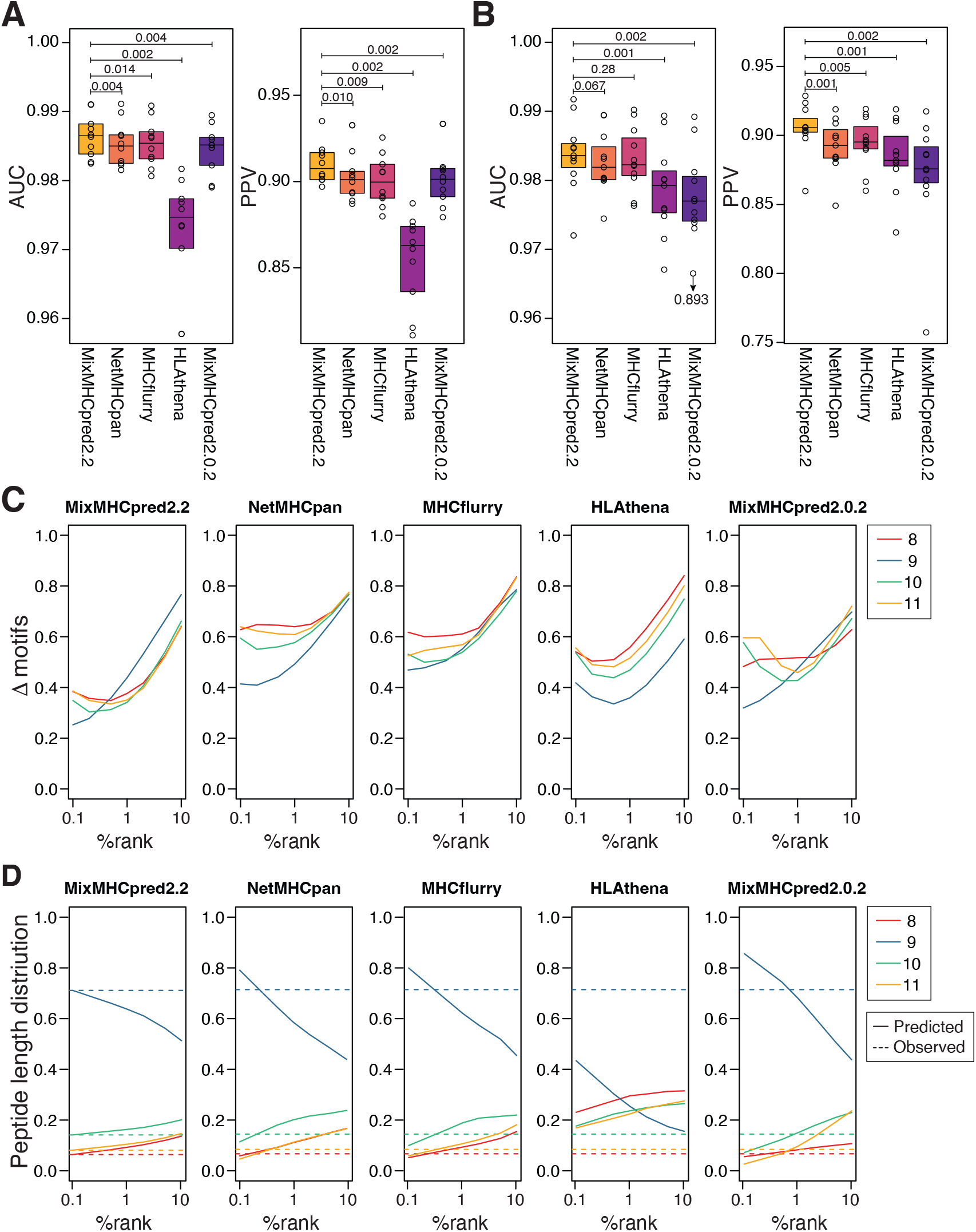
Models of HLA-I binding specificities and peptide length distributions improve predictions of naturally presented HLA-I ligands. (A) Boxplot of AUC and PPV values for the different predictors considered in this study applied on the 10 HLA-I peptidomics samples from (Gfeller et al., 2018). (B) AUC and PPV values obtained for HLA-I peptidomics samples from (Pyke et al., 2021). (C) Euclidean distance between observed and predicted HLA-I binding motifs at different %rank thresholds for each HLA-I ligand predictor (average over all alleles available in each predictor). (D) Predicted peptide length distributions at different %rank thresholds for each HLA-I ligand predictor (average over alleles available in all predictors). Dashed lines show the peptide length distributions observed in naturally presented HLA-I ligands (average over all alleles). P-values in (A) and (B) were computed with paired Wilcoxon test.

Different performance in predicting naturally presented HLA-I ligands could originate either from differences in modeling binding specificities or peptide length distributions. To explore these two aspects of HLA-I ligand predictors, we first computed the Euclidean distance between motifs predicted by each predictor at different %rank thresholds and those observed experimentally in HLA-I peptidomics data (Figure 2C, see Methods). We observed that motifs predicted with HLAthena and MixMHCpred2.0.2 (which had lower AUC and PPV in Figure 2A-B) were not more distant from those observed in HLA-I peptidomics data compared to those predicted by NetMHCpan and MHCflurry. This suggests that some of the differences observed in Figure 2A-B may come from better modeling of peptide length distributions of HLA-I molecules. To explore this hypothesis, we computed the predicted peptide length distributions at different %rank thresholds (see Methods). We then compared these predicted peptide length distributions with those observed in HLA-I peptidomics samples (Figure 2D). Overall, we observed similar values for MixMHCpred2.2, NetMHCpan4.1 and MHCflurry2.0. MixMHCpred2.0.2 displayed less stable distributions across %rank thresholds, including an over-representation of longer peptides for high %rank (i.e., %rank between 2% and 10%) and an under-representation of such peptides for small %rank (i.e., %rank<1). HLAthena displayed a very clear under-representation of 9-mers, and over-representation of 8-, 10- and 11-mers across all %rank thresholds. The discrepancy was also observed when considering peptide length distributions from mono-allelic HLA-I peptidomics data (Figure S3E). These observations suggest that integrating peptide lengths, either as separate input nodes in neural networks (NetMHCpan and MHCflurry) or by stable renormalization of the raw scores (MixMHCpred2.2), is important to accurately capture the length distribution of naturally presented HLA-I ligands across different alleles.

### Models of TCR recognition improve predictions of neo-epitopes

To expand upon previous attempts to capture biochemical properties of epitopes that increase TCR recognition propensities (Calis et al., 2013; Chowell et al., 2015; Schmidt et al., 2021), we collected data from 70 recent neo-antigen studies. This resulted in 596 verified immunogenic neo-epitopes, as well as 6084 non-immunogenic peptides tested experimentally (see Methods and Dataset S3). Most of the immunogenic and non-immunogenic peptides were previously selected based on HLA-I ligand predictors and as a result show much higher predicted binding to HLA-I compared to random peptides (Figure 3A). To correct for this bias in our data, we further included for each neoepitope 99 peptides randomly selected from the same source protein as additional negatives (see Methods and Dataset S3). We then used these data to train a PRedictor of Immunogenic Epitope (PRIME2.0). PRIME2.0 is based on a neural network and uses as input features the predicted HLA-I presentation score (-log(%rank) of MixMHCpred2.2), the amino acid frequency at positions with minimal impact on binding to HLA-I and more likely to face the TCR (Schmidt et al., 2021), and the length of the peptide (Figure 3B, see Methods). Compared to our previous work (PRIME1.0 (Schmidt et al., 2021)), the training set of PRIME2.0 is more realistic in terms of predicted HLA-I binding of the negatives (i.e., broad coverage of the range of %rank values without major enrichment in predicted ligands). We performed multiple crossvalidation based on randomly splitting the data (standard 10-fold cross-validation), iteratively excluding specific alleles (leave-one-allele-out cross-validation), or iteratively excluding data from specific studies (leave-one-study-out cross-validation) (see Methods). The last two cross-validations are important to minimize the risk that our model captures properties specific for one over-represented allele (e.g., HLA-A*02:01) or one large study that would not apply to other alleles or other studies. Overall, we observed improved predictions with PRIME2.0 (Figure 3C and Figure S4A), even if most of the neo-epitopes considered in this work had been predicted by NetMHCpan and several of them are part of the training of NetMHCpan, MHCflurry and PRIME1.0. We also restricted our benchmark to peptides experimentally tested (i.e., excluding random negatives from the test sets) (Figure 3D and Figure S4B). Again, PRIME2.0 displayed better performance than HLA-I ligand predictors. In this case PRIME1.0 performed similarly to PRIME2.0, consistent with the fact that PRIME1.0 was mainly trained on peptides (both immunogenic and non-immunogenic) with high predicted presentation on HLA-I molecules.

**Figure 3:**
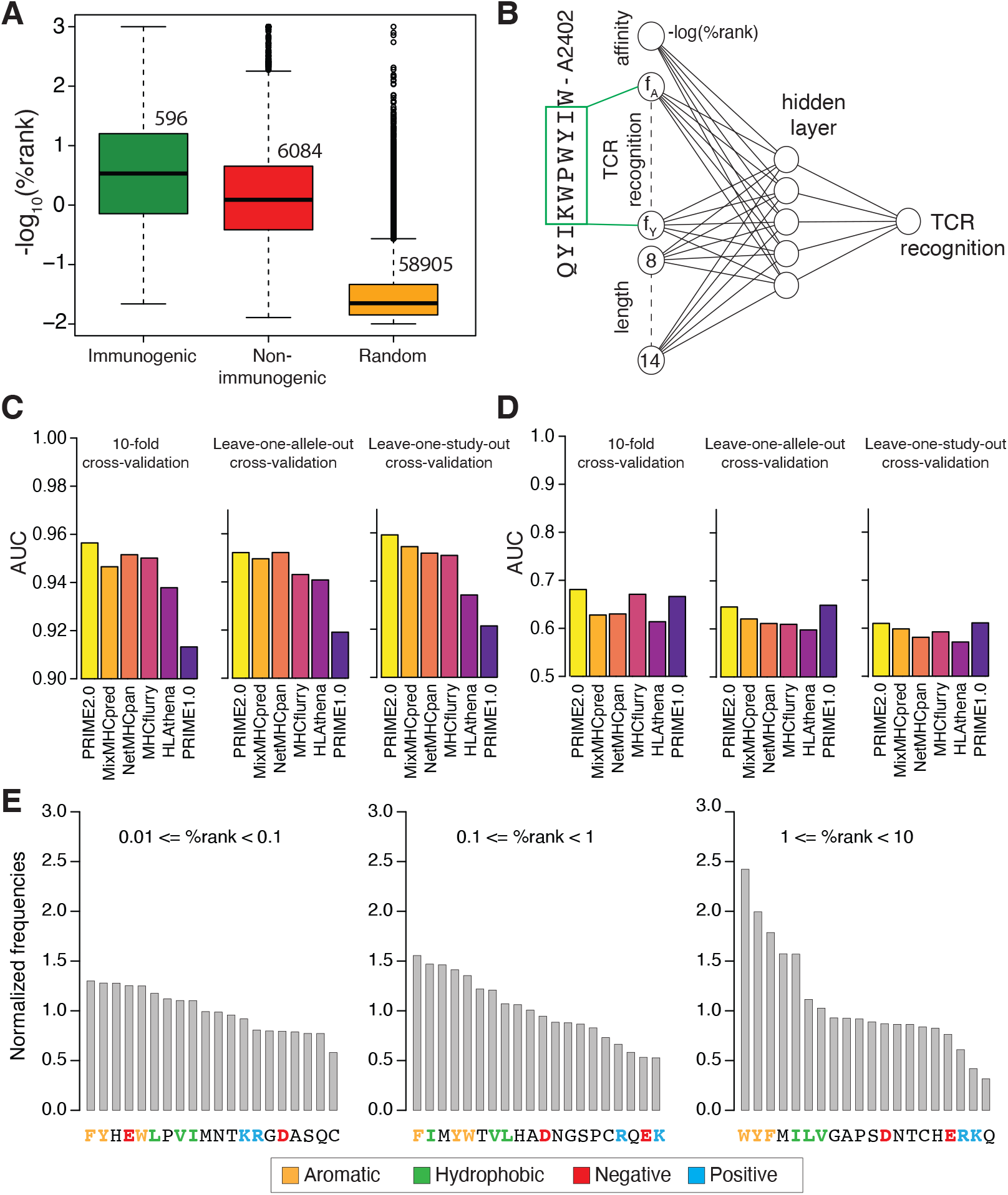
Models of TCR recognition improve predictions of neo-epitopes. (A) experimentally validated immunogenic (green) and non-immunogenic (red) peptides, as well as random peptides (orange) used to train PRIME. (B) Architecture of neural network of PRIME2.0. The first input node corresponds to the predicted binding to the HLA-I allele (-log(%rank) from MixMHCpred2.2). The next 20 nodes correspond to amino acid frequencies on residues with minimal impact on predicted affinity to the HLA-I allele (green box). These positions were determined as previously described (Schmidt et al., 2021). The last seven nodes correspond to the length of the peptide (i.e., 8 to 14, one-hot encoding). (C) Benchmarking of PRIME2.0 based on 10-fold cross-validation, leave-one-allele-out cross-validation and leave-one-study-out crossvalidation. Each bar shows the average AUC within the different types of crossvalidations (see also Figure S4A). (D) Same cross-validation as in (C) after excluding randomly generated negatives in the test set (see also Figure S4B). (E) Normalized amino acid frequencies at positions with minimal impact on predicted affinity to HLA-I for immunogenic versus non-immunogenic peptides used to train PRIME2.0 within different ranges of predicted HLA-I binding (%rank of MixMHCpred).

To further interpret the impact of amino acids found in epitope positions more likely to impact TCR recognition (green box in Figure 3B), we compared the frequency of these amino acids between the positives (i.e., epitopes) and the negatives (i.e., non-immunogenic peptides) used in the training of PRIME2.0 for different ranges of MixMHCpred %ranks (Figure 3E). Our results reveal an enrichment in aromatic and hydrophobic residues among epitopes and a depletion of charged or polar residues, which is consistent with previous studies (Calis et al., 2013; Schmidt et al., 2021). The enrichment is especially pronounced for epitopes with predicted low binding to HLA-I (%rank between 1% and 10%). These observations support the following model of TCR recognition. For high affinity HLA-I ligands the presence of specific amino acids n the region which is recognized by the TCR is less important because the high stability of the peptide-HLA-I complex increases the probability of stable TCR binding. Conversely, for low affinity HLA-I ligands, the presence of specific amino acids favoring TCR recognition becomes more important and helps counterbalancing the lower stability/affinity of the peptide-HLA-I complexes.

### Immunogenicity predictions reveal SARS-CoV-2 CD8^+^ T-cell epitopes

To further illustrate the use of PRIME, we prospectively applied it to the proteome of SARS-CoV-2 and selected a list of 213 peptides with PRIME2.0 %rank lower or equal to 0.5 with at least one of the 15 most common HLA-I alleles (see Methods and Dataset S4A-B). We then *in vitro* primed CD8^+^ T cells from 6 donors (Dataset S4C) with pools of the predicted peptides and deconvolved the IFNg ELISpot responses to the level of single epitopes (see Methods and Figure 4A). Three donors had been tested positive for SARS-CoV-2 (i.e., 1GZ0, 1HHU, 1HHT) and no donor had been vaccinated (samples collected early 2020). In total, we could identify 19 immunogenic peptides, with 2 of them (YFIASFRLF and QWNLVIGFLF) identified in two different donors (Table 1). Eight of these epitopes had not been observed in previous studies including the two identified in multiple donors (Table 1). Three additional epitopes had been reported with other allelic restrictions (LYLYALVYF, FTSDYYQLY, YFPLQSYGF). To validate these observations, we used peptide-HLA multimers to stain CD8^+^ T cells recognizing nine of these epitopes in four donors for which enough cells were available (Leu163, Leu184, 1HHT and 1HHU). All epitopes could be confirmed (Figure 4B). We then measured the functional avidity (EC_50_) of the CD8^+^ T cells recognizing these epitopes. The functional avidity displayed some variability, ranging from low micro-molar to sub-nano-molar values (Figure 4C). The highest avidity was observed for the HLA-A*29:02 restricted YFPLQSYGF epitope in a SARS-CoV-2 positive donor (1HHU). This epitope had been previously observed in patients with a restriction to HLA-A*24:02 (Saini et al., 2021). HLA-A*29:02 and HLA-A*24:02 are part of the same HLA-I supertype (A24) and display some overlap in their binding motifs, including preference for F at both P2 and PΩ anchor residues. This suggests that the YFPLQSYGF epitope may be immunogenic in several patients with HLA-I alleles of the A24 supertype.

**Figure 4:**
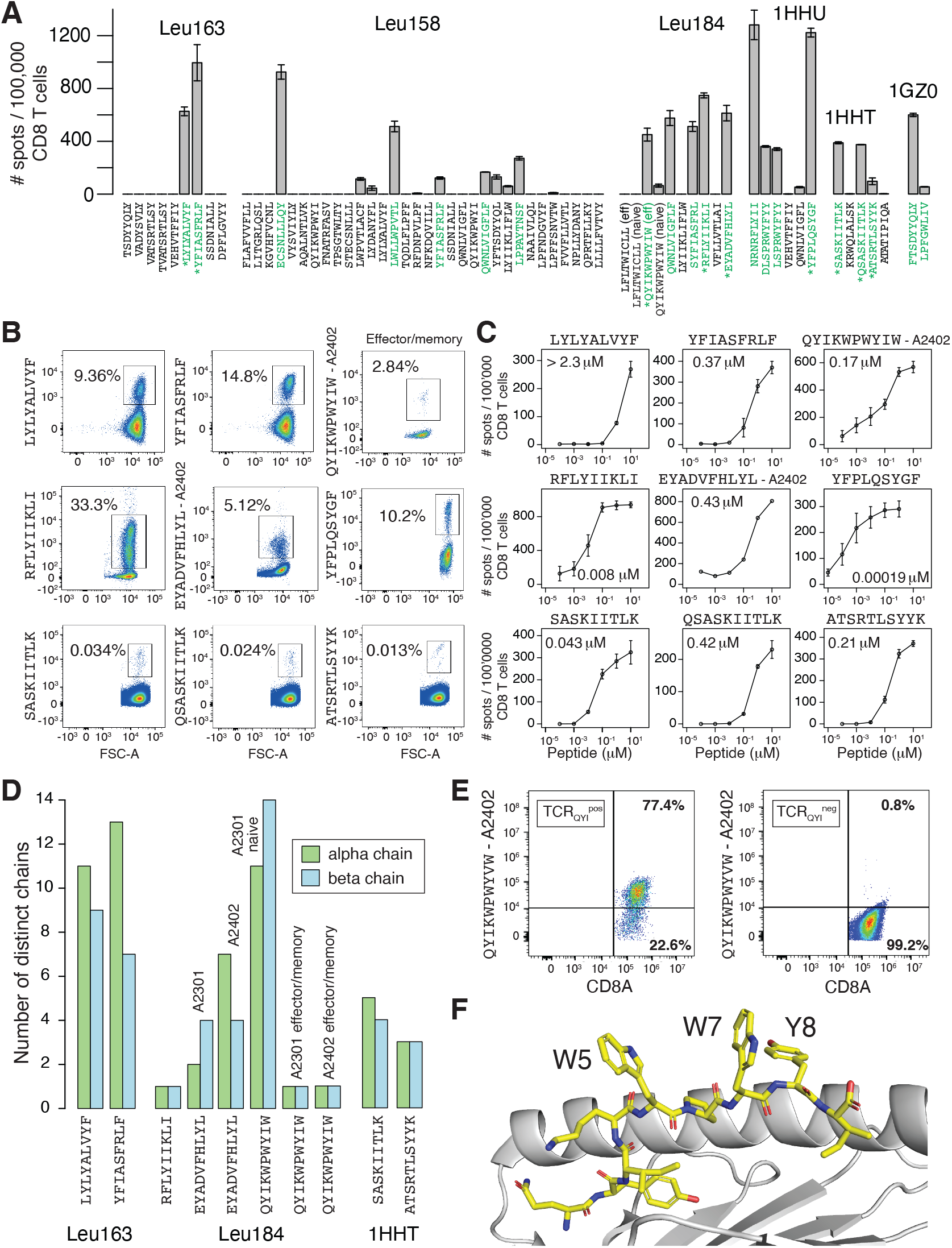
Immunogenicity predictions reveal SARS-CoV-2 CD8^+^ T-cell epitopes. (A) IFNγ ELISpot results for the peptides tested individually (i.e. after deconvolution of the pools). Immunogenic peptides are shown in green. Stars indicate peptides for which enough CD8^+^ T cells were available for peptide-HLA multimer validation and functional avidity assays. Donors are indicated above each peptide group. For donor Leu184, two epitopes (LFLTWICLL and QYIKWPWYIW) were tested with both effector/memory and naïve CD8^+^ T cells. (B) Staining of CD8^+^ T cells with peptide-HLA multimers for nine epitopes from donors for which enough CD8^+^ T cells could be obtained (i.e. donors Leu163, Leu184, 1HHT and 1HHU, see Table 1). For QYIKWPWYIW, effector/memory CD8^+^ T cells were used and the multimer was built with HLA-A*24:02 (see also Figure S5C). (C) Functional avidity (EC_50_). Error bars represent the standard deviation of two replicate, except for EYADVFHLYL where only one replicate could be performed due to limited amount of CD8^+^ T cells. (D) Number of distinct alpha and beta chains identified in TCRs recognizing the seven epitopes for which TCR sequencing could be performed. (E) QYIKWPWYVW – HLA-A*24:02 multimer staining of TCR^-^ Jurkat cells transfected with TCR_QYI_. The negative control on the right represents TCR^-^ Jurkat cells not transfected with any TCR. (F) Crystal structure (PDB: 7EJL) of the 9-mer QYIKWPWYI epitope (yellow) in complex with HLA-A*24:02 (grey). The aromatic residues at non-anchor positions (W5, W7 and Y8) point outside of the HLA-I binding pocket and towards the TCR binding interface. For clarity, the a1 helix of the HLA-I is not shown.

**Table 1:**
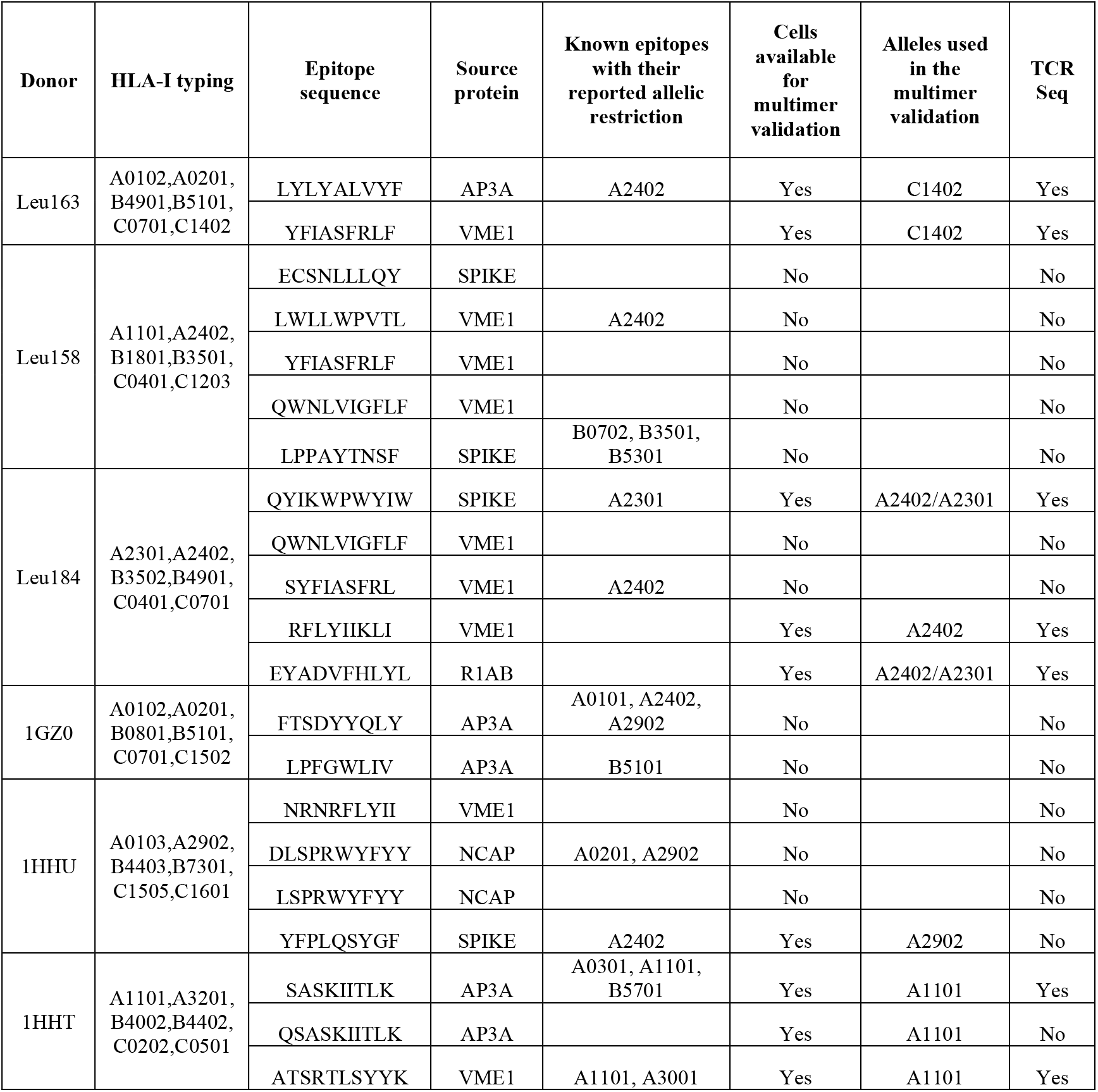
List of immunogenic SARS-CoV-2 epitopes.

To gain insights in the clonality of the CD8^+^ T cell populations recognizing these epitopes, we sorted CD8^+^ T cells recognizing seven of these epitopes and sequenced separately the alpha and beta chains of their TCRs (Figure 4D, Dataset S5). Different epitopes were recognized by different numbers of TCRs. For epitopes recognized by several TCRs, one or two alpha and beta chains had significantly higher frequency, suggesting that the recognition may be primarily driven by the pairing of such chains (Figure S5A). For the QYIKWPWYIW epitope from the Spike protein (donor Leu184), naïve and effector/memory CD8^+^ T cells recognizing this epitope were sorted separately (Figure S5B). We observed a high diversity of TCR chains among naïve CD8^+^ T cells (Figure 4D). Reversely, a unique clone (TCR_QYI_: TRAV20*01-CAALNYGGATNKLIF-TRAJ32*01 and TRBV4-3*01-CASSPSGGAYEQYF-TRBJ2-7*01) was found in the effector/memory CD8^+^ T cells (Figure 4D). This unique TCR was identified in effector/memory CD8^+^ T cells recognizing QYIKWPWYIW displayed both on HLA-A*23:01 and HLA-A*24:02 (Figure S5C, Dataset S5), as expected since HLA-A*23:01 and HLA-A*24:02 have very high sequence similarity and almost identical binding motifs (Figure S1). We next asked if the same TCR could be found in other individuals. The same beta chain was found in 19 SARS-CoV-2^+^donors in the ImmuneCODE database, which is a large repertoire of TCRβ chains from SARS-CoV-2^+^ donors (Nolan et al., 2020). The same alpha and beta chains were also found in the TCR repertoire of one of the two SARS-CoV-2^+^ donors analyzed in a recent study (Minervina et al., 2021). Moreover, the same alpha chain and a highly similar beta chain (same CDR3β sequence) were found in the other donor of this study (Figure S5D). Both donors were HLA-A*24:02^+^. These observations suggest that the recognition of the QYIKWPWYIW epitope may be mediated by the same TCR in multiple donors.

The donor where recognition of the QYIKWPWYIW epitope was observed (Leu184) had not been tested positive for SARS-CoV-2 and had not received any SARS-CoV-2 vaccine. Therefore we hypothesized that the monoclonal population of effector/memory CD8^+^ T cells recognizing this epitope could originate from previous exposure to other coronaviruses. This hypothesis is supported by the fact that the QYIKWPWYIW epitope is almost perfectly conserved in the Spike protein of SARS-CoV-1 (QYIKWPWYVW, I>V mutation at position 9). To verify our hypothesis, we stained cells transfected with TCR_QYI_ with the QYIKWPWYVW – HLA-A*24:02 multimer. Our results demonstrate that TCR_QYI_ is able to recognize this SARS-CoV-1 epitope (Figure 4E). These results are consistent with the observation that previous exposure to other coronaviruses can confer some immunity to SARS-CoV-2 (Braun et al., 2020; Loyal et al., 2021).

In a recent study, the 9-mer peptide (QYIKWPWYI) fully overlapping with the 10-mer epitope recognized by TCR_QYI_ was shown to elicit an immuno-dominant CD8^+^ T-cell response and the QYIKWPWYI – HLA-A*24:02 complex was crystallized (Shimizu et al., 2021). This structure shows that the three non-anchor aromatic sidechains shared with the 10-mer investigated in our work (i.e., W5, W7 and Y8) are all facing outside of the HLA-I binding site and therefore are likely to interact with the TCR (Figure 4F). The presence and orientation of the aromatic sidechains in this immuno-dominant epitope of the Spike protein are consistent with the model of improved TCR recognition propensity of aromatic residues, which underlies the PRIME algorithm.

## Discussion

CD8^+^ T-cell epitopes play central roles in immune responses against infectious diseases and cancer, and represent promising targets for personalized cancer immunotherapy treatments. In this work, we trained both a predictor of antigen presentation (MixMHCpred) and a predictor of immunogenicity (PRIME). By expanding the training set and optimizing the algorithms, we could demonstrate improved predictions for both HLA-I ligands and class I neo-epitopes.

A key element of any machine learning predictor is the quality and depth of the training data. Consistent with previous studies (Alvarez et al., 2019; Fritsche et al., 2021; Gfeller et al., 2018), our results reveal that different types of putative contaminants can be found in both poly- and mono-allelic HLA-I peptidomics data. Contaminants include peptides with trypsin-like motifs or peptides coming from other HLA-I alleles, including in samples that were assumed to be mono-allelic. These results emphasize the importance of carefully applying quality controls before using such data for training HLA-I ligand predictors (Fritsche et al., 2021).

The comparison between observed and predicted HLA-I binding motifs demonstrated lower distances for motifs predicted by MixMHCpred2.2 compared to other predictors (Figure 2C). While this may contribute to the improved predictions observed in the independent validations of Figure 2A-B, it is important to point out that this analysis is influenced by the fact that MixMHCpred2.2 was trained on the same data used to derive the observed motifs. Results in Figure 2C should therefore not be interpreted as validation of the improved predictions of MixMHC2pred2.2, but only as an indication that other tools do not differ much in the modelling of HLA-I binding motifs, and that the most important differences are found at the level of peptide length distributions.

The analysis of the data used to train the predictor of immunogenicity (PRIME) confirmed the importance of aromatic residues, especially tryptophan. In line with previous studies and crystal structures of TCR-peptide-MHC complexes (Calis et al., 2013; Devlin et al., 2020; Schmidt et al., 2021; Shimizu et al., 2021), we suggest that this preference reflects the ability of tryptophan to engage into stable molecular interactions with the TCR. However, we cannot exclude that other factors play a role in the importance given to tryptophan in PRIME. First, tryptophan tends to be slightly depleted in MS-based HLA-I peptidomics studies (Abelin et al., 2017; Bassani-Sternberg et al., 2017). This may bias HLA-I ligand predictors trained on such data, and PRIME may be correcting for this bias. Second, recent studies have demonstrated that peptides genomically encoded with a W can undergo a W>F substitution during protein synthesis (Pataskar et al., 2022). Considering that several studies used tandem mini-genes to identify neo-epitopes, we cannot exclude that some of the W-containing epitopes used in the training of PRIME were actually presented on HLA-I molecules with a W>F substitution, which contributed to overcome central tolerance and increase their immunogenicity. This supports a model where tryptophan containing protein segments are especially promising neo-epitope candidates, both in terms of improving TCR recognition and overcoming central tolerance.

Applying our tool to the SARS-CoV-2 proteome, we could uncover several epitopes, including one (QYIKWPWYIW) recognized by a monoclonal population of antigen-experienced CD8^+^ T cells with an effector/memory phenotype. This epitope has very high homology with SARS-CoV-1 (QYIKWPWYVW) and is 100% conserved in all common variants of SARS-CoV-2. This suggests that CD8^+^ T-cell responses elicited against this epitope by previous infection, vaccination or cross-reactivity with SARS-CoV-1 may be effective across all SARS-CoV-2 variants.

Overall, our work provides improved approaches for both antigen presentation and TCR recognition predictions of class I epitopes, and reveals new SARS-CoV-2 epitopes. In terms of HLA-I ligand predictions, a decent accuracy had already been reached by many existing tools (Gfeller et al., 2018; O’Donnell et al., 2020; Reynisson et al., 2020). Much harder is the task of predicting TCR recognition, both because of the smaller size of the training data and because of the many other factors that influence TCR recognition (e.g., co-receptors, cytokines, etc.). Efforts focusing on generating high quality immunogenicity training data and developing machine learning frameworks to harness these data will be key to further improve class I epitope predictions.

## Methods

### Data collection and curation

Naturally presented HLA-I ligands of length 8 to 14 were collected from 244 samples, coming from 24 different HLA-I peptidomics studies (see Dataset S1). These comprise both mono- and poly-allelic samples. All data were retrieved from the original studies to prevent any filtering based on HLA-I ligand predictors. When only filtered data had been published in the original studies, access to unfiltered data was kindly provided to us by the authors of these studies. All samples were processed with the motif deconvolution algorithm MixMHCp in order to identify shared HLA-I motifs across samples sharing the same alleles, following our previously established procedure (Bassani-Sternberg and Gfeller, 2016; Bassani-Sternberg et al., 2017; Gfeller et al., 2018). All samples were manually reviewed and peptides assigned to motifs that could not be unambiguously assigned to one allele were not considered. Peptides assigned to the flat motif (trash) in MixMHCp or to motifs corresponding to alleles not supposed to be in the sample were considered as predicted contaminants. The final dataset of naturally presented HLA-I ligands consists of 258,814 unique peptides, representing 384,070 peptide-HLA-I interactions with 119 different HLA-I alleles. 59 alleles were observed in both mono- and poly-allelic samples, 22 only in poly-allelic samples, and 38 only in mono-allelic samples. This curated set of naturally presented HLA-I ligands is available in Dataset S2. Binding motifs of HLA-I alleles were plotted with ggseqlogo (Wagih, 2017).

Immunogenic neo-epitopes were retrieved from several neo-antigen studies and were completed by neo-epitope data from IEDB (Vita et al., 2019) (tcell_full_v3.csv file, downloaded on March 27, 2021), excluding potential overlap (see Dataset S3). Both immunogenic and non-immunogenic mutated peptides were considered. This resulted in 596 experimentally verified neo-epitopes (10 8-mers, 391 9-mers, 148 10-mers and 47 11-mers) and 6084 experimentally verified non-immunogenic peptides (Dataset S3).

### Computing peptide length distributions

Peptide length distributions were established by computing the fraction of naturally presented HLA-I ligands of each length *l* ∈ [8,14] for each allele. For each of the 59 alleles found in both mono-allelic and poly-allelic samples, peptide length distributions were also computed separately for ligands coming from mono-allelic and poly-allelic samples (Figure 1C).

### Training of MixMHCpred

MixMHCpred2.2 was trained based on our curated set of naturally presented HLA-I ligands, following the procedure described in (Gfeller et al., 2018). The main difference consists of a more stable modelling of peptide length distributions. In mathematical terms, the score of a peptides *X* of length *L* with allele *h* is given by:

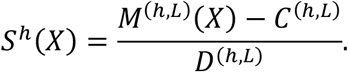

*M*^(*h,L*)^(*X*) represents the raw score of peptide *X* given by the Position Weight Matrices representing the motif of allele *h* for *L*-mers, including normalization by background frequencies and BLOSUM62 based pseudocounts, as described in (Gfeller et al., 2018). The correction factors *D*^(*h,L*)^ were computed so that *S^h^*(*X*) has a standard deviation of 1 over a set of 100’000 peptides of length *L* randomly selected from the human proteome (i.e., *D*^(*h,L*)^ represents the standard deviation of the scores of these peptides). The correction factors *C*^(*h,L*)^ were computed so that the length distribution of the top 0.1% of 700’000 random peptides (taken from the human proteome with uniform length distribution between 8- and 14-mers) follows exactly the peptide length distribution of allele *h* observed in HLA-I peptidomics data. In mathematical terms, defining *P^h^*(*L*) as the experimental peptide length distribution for a given allele *h*, *C*^(*h,L*)^ corresponds to the raw score 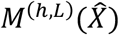, where 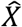 represents the *L*-mer peptide ranked 100’000 × 0.001 × (*L*^max^ − *L*^min^ + 1) × *P^h^*(*L*) among the set of 100’000 random *L*-mer peptides, with *L*^min^ = 8 and *L*^max^ = 14. Given the observed discrepancies between peptide length distributions from mono and poly-allelic samples (Figure 1C), peptide length distributions from poly-allelic samples were always used, when available. %ranks given as output of MixMHCpred2.2 were estimated based on the distribution of scores *S^h^*(*X*) of a set of 700’000 random peptides (100’000 of each length from 8 to 14), as done in other HLA-I ligand predictors. Consistent with recommendations for other tools (Reynisson et al., 2020), these %rank should be used for ranking candidates to be experimentally validated. The new version of MixMHCpred was benchmarked against NetMHCpan4.1 (Reynisson et al., 2020), MHCflurry2.0 (O’Donnell et al., 2020), HLAthena (Sarkizova et al., 2019) and MixMHCpred2.0.2 (Gfeller et al., 2018) using naturally presented HLA-I ligands identified in unmodified tissues. To ensure that the HLA-I peptidomics samples used for this benchmark were not part of the training of any of these tools, we used (i) HLA-I peptidomics datasets coming from 10 menigioma samples measured in (Gfeller et al., 2018) that were not integrated in the training of any version of MixMHCpred, NetMHCpan, MHCflurry or HLAthena and were not uploaded in IEDB and (ii) HLA-I peptidomics samples from (Pyke et al., 2021) which were published after the latest release of these tools, excluding sample ‘ 1180157F’ due to ambiguity in HLA-I typing. 4-fold excess of random negatives were added by randomly selecting peptides from the human proteome. For this comparison, we restricted to peptides of length 8 to 11, since HLAthena cannot be run for longer peptides. PPV in the top 20% (which is equivalent to recall with 4-fold excess of random negatives) and AUC values were computed for each sample and each predictor.

### Comparing predicted and experimental HLA-I motifs

To compare HLA-I motifs predicted by different tools (Figure 2C), we randomly selected 100’000 peptides from the human proteome for each length *l* ∈ [8,11] (higher values of *l* were not considered in this comparison, since the number of ligands is much smaller, and HLAthena is limited to 8- to 11-mers). For each allele and each HLA-I predictor, we computed the %rank of each peptide. Predicted motifs for different %rank thresholds were computed based on the peptides that had a %rank lower or equal to the thresholds, by computing the amino acid frequencies at each positions (20 x *l* matrices). These frequencies were compared to those observed in experimental HLA-I ligands, using the Euclidean distance (Δ motifs). The average of the motif distances was plotted in Figure 2C for each peptide length and each %rank threshold. Only cases with at least 20 ligands predicted by all predictors and observed experimentally were considered.

### Comparing predicted and experimental peptide length distributions

As with predicted motifs, peptide length distributions predicted by each predictor at different %rank thresholds were computed based on 100’000 randomly selected peptides from the human proteome for each length *l* ∈ [8,11]. The average peptide length distribution over all alleles available in all predictors is shown in Figure 2D for each predictor and each %rank threshold.

### Training of PRIME

The new version of PRIME (v2.0) was trained using a fully connected neural network with 5 hidden nodes (*mlp* package in R (Bergmeir and Benítez, 2012)). The input layer consists of 28 nodes (Figure 3B). The first input node encodes the predicted binding to the HLA-I molecule (-log(%rank), predicted by MixMHCpred). The twenty next input nodes encode amino acid frequencies at positions with minimal impact on predicted affinity to HLA-I and more likely to interact with the TCR (Schmidt et al., 2021). The last seven input nodes encode the length of the peptide (one-hot encoding of lengths 8 to 14).

The set of experimentally verified immunogenic and non-immunogenic peptides was used to train PRIME2.0. As this set of peptides is heavily skewed towards peptides with high predicted affinity (Figure 3A), 99-fold excess of negatives were further added by randomly selecting for each immunogenic neo-epitope 99 peptides from the same source protein (non-mutated), for a total of 58,905 random peptides (for one neoepitope, the source protein could not be found, and no random peptide was included for this neo-epitope). The length of these negatives was randomly chosen between 8 and 14. The use of only human (mutated) peptides in both positives and negatives prevents potential biases in amino acid frequencies due to different GC content across different organisms.

To benchmark the new version of PRIME, we first performed a standard 10-fold cross-validation, by randomly splitting the data in ten groups, iteratively training the model on nine groups and testing on the remaining one. Given that our dataset of immunogenic neo-epitopes is skewed towards frequent HLA-I alleles and towards studies where many neo-epitopes had been reported, we also performed a leave-one-allele-out, respectively a leave-one-study-out, cross-validation, using iteratively as test set each allele, respectively each study, with more than two experimentally validated immunogenic and two experimentally validated non-immunogenic peptides.

### Predictions of SARS-CoV-2 epitopes

The SARS-CoV-2 reference proteome was downloaded from UniProt on March 22, 2020 and peptides of length 8 to 11 were retrieved. The list of HLA-I alleles was established by taking the top 15 most frequent alleles in the TCGA cohort (Dataset S4B). Peptides with a %rank lower or equal to 0.5 for PRIME2.0 for at least one allele and coming from the five proteins SPIKE, VME1, VEMP, NCAP, AP3A were first considered. A few peptides from R1AB were further manually included as they came from regions with several predicted epitopes for multiple alleles. The final list consists of 213 peptides (Dataset S4A).

### Donors and regulatory issues

Six donors were recruited (Leu163, Leu158, Leu184, 1GZ0, 1HHT, 1HHU). The HLA-I typing was known for all six donors and the last three donors (1GZ0, 1HHT, 1HHU) had been tested positive for SARS-CoV-2 (PCR tests) (Dataset S4C). The recruitment and blood withdrawal were approved by regulatory authorities and all donors signed informed consents (protocol TRP0014, BASEC_ID : 2018-01838).

### Identification of SARS-CoV-2 epitopes

The 213 peptides were purchased at ThermoFisher (>80 % purity), solubilized in DMSO at 10 mM, aliquoted and kept at −80°C. CD8^+^ T cells were isolated (ref 130-045-201, Miltenyi) from cryopreserved PBMC (for SARS-CoV-2 positive donors) or fresh leukapheresis (for SARS-CoV-2 negative donors). For SARS-CoV-2 positive donors (1HHU, 1HHT, 1GZ0), due to the limited number of PBMCs, total CD8^+^ T cells were used for further *in vitro* stimulation. For the other three donors (Leu163, Leu158. Leu184), naive and effector/memory CD8^+^ T cells were isolated by Fluorescence-activated Cell Sorting (FACS) upon staining with anti-CD8 antibody (344710 BioLegend), anti-CCR7 antibody (353227 BioLegend) and anti-CD45RA antibody (304108 BioLegend) for 30 min at 4°C. After three washes with FACS buffer (PBS 0.5 % FBS 2 mM EDTA) cells were incubated 10 min with DAPI (Sigma 10236276001) at 250 nM and washed again three times. Total CD8^+^ T cells (donors 1GZ0, 1HHT, 1HHU), naïve (CCR7^+^ and CD45RA^+^) CD8^+^ T cells (donors Leu163, Leu158, Leu184) and effector/memory (CD45RA^-^) CD8^+^ T cells (donor Leu184 – not enough effector/memory cells available for the other donors) were collected separately and then co-incubated (10^6^ mL^-1^) with autologous irradiated CD8-depleted PBMCs and pools of 11 to 24 peptides (1 μM) in RPMI supplemented with 8 % human serum and IL-2 (50 IU mL^-1^ for 48 h and then switch 1mL of media with 150 IU mL^-1^ every 48h, split as necessary to get minimum 10^6^ Cell.mL^-1^). IFNγ Enzyme-Linked ImmunoSpot (ELISpot) was performed at day 12 post-stimulation. One day before ELIspot, cells were incubated in r8 media without IL2. ELISpot assays were performed using precoated 96-well ELISpot plates (Mabtech 3420-2APT-10) and counted with Bioreader-6000-E (BioSys). All peptide pools giving a specific response (considered if at least 10 spots for 100 000 incubated cells and 2 times the background signal, obtained by incubation of cells without peptide) were deconvoluted by repeating ELISpot assays with individual peptides.

### Peptide-HLA multimer validation of SARS-CoV-2 epitopes and sorting of CD8^+^ T cells

Peptides found as immunogenic in the ELISpot assays were resynthesized with a purity >95 % and used for production of peptide-HLA multimers (Peptide and Tetramer Core Facility of the University Hospital of Lausanne). CD8^+^ T cells were incubated with multimers (1/50 dilution) 45 min at 4°C in FACS buffer (PBS supplemented with 0.5 % FBS and 2 mM EDTA), isolated by FACS and either directly used for TCR sequencing or expanded with autologous irradiated CD8-depleted feeders in RPMI supplemented with 8% human serum, phytohemagglutinin (1 μg mL^-1^) and IL2 (150 IU mL^-1^).

### Functional avidity assay

Functional avidity of antigen-specific CD8^+^ T-cell responses was assessed by performing *in vitro* IFNγ Enzyme-Linked ImmunoSpot (Mabtech) assay with limiting peptide dilutions (ranging from 10 μg mL^-1^ to 10 pg mL^-1^) as described earlier (Viganò et al., 2012). For all peptide concentrations, ELISpot signals were measured in two replicates and the average of the two replicates was used to compute EC_50_ values. EC_50_ values reported in Figure 4C were computed by fitting sigmoid curves with the “ec50estimator” package in R (https://github.com/AlvesKS/ec50estimator). For EYADVFHLYL, enough cells were available for only one replicate. For the HLA-A*29:02 restricted YFPLQSYGF epitope, single clones were isolated and the EC_50_ values represent the average over all clones coming from two different pools (error bars represent the standard deviation between the average values in the two pools). For this epitope, peptide concentrations ranging from 10^-11^ to 10^-6^ were used, as the first response was stronger than for other epitopes (Figure 4C).

### Bulk TCR sequencing

mRNA was extracted using the Dynabeads mRNA DIRECT purification kit according to the manufacturer instructions (ThermoFisher) and was then amplified using the MessageAmp II aRNA Amplification Kit (Ambion) with the following modifications: *in vitro* transcription was performed at 37°C for 16 h. First strand cDNA was synthesized using the Superscript III (Thermofisher) and a collection of TRAV/TRBV specific primers. Unique Molecular identifiers (UMI) of length 9 were added to each read. TCRs were then amplified by PCR (20 cycles with the Phusion from NEB) with a single primer pair binding to the constant region and the adapter linked to the TRAV/TRBV primers added during the reverse transcription. A second round of PCR (25 cycles with the Phusion from NEB) was performed to add the Illumina adapters containing the different indexes. The TCR products were purified with AMPure XP beads (Beckman Coulter), quantified and loaded on the MiSeq instrument (Illumina) for deep sequencing of the TCRα/TCRβ chain.

### TCR sequence analyses

The fastq files were processed with MIGEC (Shugay et al., 2014), using default parameters to demultiplex them and identify the TCRα and TCRβ clonotypes. For each sample, the frequency of each TCR chain was computed based on UMI corrected counts. Only TCRs with more than one UMI count and representing more than 1% of the total UMI counts were considered (Dataset S5). TCRs with the same amino acid sequences were merged in Figure 4D and Figure S5A.

The beta chain of the TCR_QYI_ (TRBV4-3*01-CASSPSGGAYEQYF-TRBJ2-7*01) recognizing the QYIKWPWYIW epitope and found in effector/memory CD8^+^ T cells of Leu184 was used to search TCRβ repertoires in the ImmunoCode database (Nolan et al., 2020) through the iReceptor web platform (Corrie et al., 2018). Both the alpha and beta chains were used to query separately the TCRα and TCRβ repertoires of the two SARS-CoV-2+ patients in Minervina et al. (Minervina et al., 2021). The closest hits (Figure S5D) were defined as those having the same CDR3 sequence and the most similar CDR1 and CDR2, based on sequence identity (100% identity for the alpha chain in both donors, 100% identity for the beta chain in donor M).

### TCR_QYI_ transfection in Jurkat cells and recognition of the SARS-CoV-1 homologous epitope

TCR_QYI_ full-length a and β chains were in silico designed and obtained by Thermo Fisher Scientific as strings. Strings have been amplified and purified by silica membrane columns (NucleoSpin PCR Clean-up, Macherey-Nagel) and used as individual templates for mRNA in vitro transcription using the HiScribe T7 In vitro transcription kit (NEB), followed by lithium chloride precipitation, as instructed by the manufacturer. RNA polyadenylation and molecular size were assessed by gel electrophoresis in denaturing conditions. Purified RNA was quantified using a Qubit BR Assay kit (Thermo Fisher Scientific) and resuspended in H_2_O at 1-2μg/mL followed by storage at −80°C, until used.

TCRα and TCRβ pairs were transfected into a recipient Jurkat cell line (T cell activation bioassay NFAT, Promega) that was further engineered by knocking out the endogenous TCRα and TCRβ chains using CRISPR/Cas9 and by stable transduction with CD8A and CD8B. Cells were propagated following the manufacturer’s instructions. For TCR transfection, 1×10^6^ Jurkat cells were co-electroporated with 3 μg of each TCR chain using a Neon Transfection System 100μl kit (Thermo Fisher Scientific) with the following parameters: 1325V, 10ms, 3 pulses. After electroporation, cells were immediately resuspended in complete medium and incubated at 37°C for 18-20 hours before staining. TCR_QYI_ electroporated Jurkat cells were stained with a PE conjugated QYIKWPWYVW-HLA–A*24:02 multimer (synthesized in house), washed once and further stained with anti-CD3 APC-Fire (Biolegend), -CD4 PE-CF594, and -CD8 FITC (BD Biosciences) fluorophore-conjugated anti-human antibodies. Aqua live dye (Thermo Fisher Scientific) was used to access viability. As control, Jurkat cells electroporated with H_2_O (mock) were stained in parallel. The samples were acquired by a iQue Screener PLUS (Intellicyt) flow cytometer and analysed by FlowJoX.

## Data accessibility

MixMHCpred (v2.2) and PRIME (v2.0) are available at https://github.com/GfellerLab/ and through the web interface http://prime.gfellerlab.org/. TCR sequencing data were deposited at GEO (accession number: GSE201212). The HLA-I ligand and neo-epitope datasets used to train MixMHCpred and PRIME are available in Dataset S2 and S3.

## Acknowledgements

D.G and A.H. acknowledge support from the Swiss National Science Foundation Sinergia grant (CRSII5_193749). D.G. acknowledges support from the Swiss Cancer Research Foundation (KFS-4104-02-2017). A.H. acknowledges support from the Swiss National Science Foundation (310030_182384). G.C. is supported by the Marie-Curie fellowship (H2020-MSCA-IF-2020, No 101027973).

## Author contributions

D.G. designed the study. D.G. performed the bioinformatics analyses. D.G. and J.R. developed the bioinformatics tools. D.G. and G.C performed the TCR sequence analyses. J.S., S.B., R.G., P.G., L.Q., J.C. performed the experiments. A.H. supervised the experiments. D.G wrote the manuscript. G.C., J.S., S.B., R.G., J.R. and A.H. provided materials and feedback on the manuscript.

## Declaration of Interest

The authors declare no competing interests.

## Notes

### Competing Interest Statement

The authors have declared no competing interest.

## References

Abelin, J.G., Keskin, D.B., Sarkizova, S., Hartigan, C.R., Zhang, W., Sidney, J., Stevens, J., Lane, W., Zhang, G.L., Eisenhaure, T.M., et al. (2017). Mass Spectrometry Profiling of HLA-Associated Peptidomes in Mono-allelic Cells Enables More Accurate Epitope Prediction. Immunity 46, 315–326. https://doi.org/10.1016/j.immuni.2017.02.007.

Alvarez, B., Reynisson, B., Barra, C., Buus, S., Ternette, N., Connelley, T., Andreatta, M., and Nielsen, M. (2019). NNAlign_MA; MHC Peptidome Deconvolution for Accurate MHC Binding Motif Characterization and Improved T-cell Epitope Predictions. Mol. Cell Proteomics 18, 2459–2477. https://doi.org/10.1074/mcp.TIR119.001658.

Andreatta, M., Alvarez, B., and Nielsen, M. (2017). GibbsCluster: unsupervised clustering and alignment of peptide sequences. Nucleic Acids Res. 45, W458–W463. https://doi.org/10.1093/nar/gkx248.

Balachandran, V.P., Łuksza, M., Zhao, J.N., Makarov, V., Moral, J.A., Remark, R., Herbst, B., Askan, G., Bhanot, U., Senbabaoglu, Y., et al. (2017). Identification of unique neoantigen qualities in long-term survivors of pancreatic cancer. Nature 551, 512–516. https://doi.org/10.1038/nature24462.

Bassani-Sternberg, M., and Gfeller, D. (2016). Unsupervised HLA Peptidome Deconvolution Improves Ligand Prediction Accuracy and Predicts Cooperative Effects in Peptide-HLA Interactions. J. Immunol. 197, 2492–2499. https://doi.org/10.4049/jimmunol.1600808.

Bassani-Sternberg, M., Chong, C., Guillaume, P., Solleder, M., Pak, H., Gannon, P.O., Kandalaft, L.E., Coukos, G., and Gfeller, D. (2017). Deciphering HLA-I motifs across HLA peptidomes improves neo-antigen predictions and identifies allostery regulating HLA specificity. PLoS Comput. Biol. 13, e1005725. https://doi.org/10.1371/journal.pcbi.1005725.

Bergmeir, C., and Benítez, J.M. (2012). Neural Networks in R Using the Stuttgart Neural Network Simulator: RSNNS. Journal of Statistical Software 46, 1–26..

Braun, J., Loyal, L., Frentsch, M., Wendisch, D., Georg, P., Kurth, F., Hippenstiel, S., Dingeldey, M., Kruse, B., Fauchere, F., et al. (2020). SARS-CoV-2-reactive T cells in healthy donors and patients with COVID-19. Nature 587, 270–274. https://doi.org/10.1038/s41586-020-2598-9.

Calis, J.J.A., Maybeno, M., Greenbaum, J.A., Weiskopf, D., De Silva, A.D., Sette, A., Keşmir, C., and Peters, B. (2013). Properties of MHC class I presented peptides that enhance immunogenicity. PLoS Computational Biology 9, e1003266..

Chowell, D., Krishna, S., Becker, P.D., Cocita, C., Shu, J., Tan, X., Greenberg, P.D., Klavinskis, L.S., Blattman, J.N., and Anderson, K.S. (2015). TCR contact residue hydrophobicity is a hallmark of immunogenic CD8+ T cell epitopes. Proc. Natl. Acad. Sci. U.S.A. 112, E1754–1762. https://doi.org/10.1073/pnas.1500973112.

Corrie, B.D., Marthandan, N., Zimonja, B., Jaglale, J., Zhou, Y., Barr, E., Knoetze, N., Breden, F.M.W., Christley, S., Scott, J.K., et al. (2018). iReceptor: A platform for querying and analyzing antibody/B-cell and T-cell receptor repertoire data across federated repositories. Immunol Rev 284, 24–41. https://doi.org/10.1111/imr.12666.

Devlin, J.R., Alonso, J.A., Ayres, C.M., Keller, G.L.J., Bobisse, S., Vander Kooi, C.W., Coukos, G., Gfeller, D., Harari, A., and Baker, B.M. (2020). Structural dissimilarity from self drives neoepitope escape from immune tolerance. Nat Chem Biol https://doi.org/10.1038/s41589-020-0610-1.

Di Marco, M., Schuster, H., Backert, L., Ghosh, M., Rammensee, H.-G., and Stevanović, S. (2017). Unveiling the Peptide Motifs of HLA-C and HLA-G from Naturally Presented Peptides and Generation of Binding Prediction Matrices. J. Immunol. 199, 2639–2651. https://doi.org/10.4049/jimmunol.1700938.

Duan, F., Duitama, J., Al Seesi, S., Ayres, C.M., Corcelli, S.A., Pawashe, A.P., Blanchard, T., McMahon, D., Sidney, J., Sette, A., et al. (2014). Genomic and bioinformatic profiling of mutational neoepitopes reveals new rules to predict anticancer immunogenicity. The Journal of Experimental Medicine 211, 2231–2248..

Fritsche, J., Kowalewski, D.J., Backert, L., Gwinner, F., Dorner, S., Priemer, M., Tsou, C.-C., Hoffgaard, F., Römer, M., Schuster, H., et al. (2021). Pitfalls in HLA ligandomics - How to catch a li(e)gand. Mol Cell Proteomics 100110. https://doi.org/10.1016/j.mcpro.2021.100110.

Gfeller, D., and Bassani-Sternberg, M. (2018). Predicting Antigen Presentation-What Could We Learn From a Million Peptides? Front Immunol 9, 1716. https://doi.org/10.3389/fimmu.2018.01716.

Gfeller, D., Guillaume, P., Michaux, J., Pak, H.-S., Daniel, R.T., Racle, J., Coukos, G., and Bassani-Sternberg, M. (2018). The Length Distribution and Multiple Specificity of Naturally Presented HLA-I Ligands. J. Immunol. 201, 3705–3716. https://doi.org/10.4049/jimmunol.1800914.

Koşaloğlu-Yalçin, Z., Lee, J., Greenbaum, J., Schoenberger, S.P., Miller, A., Kim, Y.J., Sette, A., Nielsen, M., and Peters, B. (2022). Combined assessment of MHC binding and antigen abundance improves T cell epitope predictions. IScience 25, 103850. https://doi.org/10.1016/j.isci.2022.103850.

Loyal, L., Braun, J., Henze, L., Kruse, B., Dingeldey, M., Reimer, U., Kern, F., Schwarz, T., Mangold, M., Unger, C., et al. (2021). Cross-reactive CD4+ T cells enhance SARS-CoV-2 immune responses upon infection and vaccination. Science 374, eabh1823. https://doi.org/10.1126/science.abh1823.

Łuksza, M., Riaz, N., Makarov, V., Balachandran, V.P., Hellmann, M.D., Solovyov, A., Rizvi, N.A., Merghoub, T., Levine, A.J., Chan, T.A., et al. (2017). A neoantigen fitness model predicts tumour response to checkpoint blockade immunotherapy. Nature 551, 517–520. https://doi.org/10.1038/nature24473.

Minervina, A.A., Komech, E.A., Titov, A., Bensouda Koraichi, M., Rosati, E., Mamedov, I.Z., Franke, A., Efimov, G.A., Chudakov, D.M., Mora, T., et al. (2021). Longitudinal high-throughput TCR repertoire profiling reveals the dynamics of T-cell memory formation after mild COVID-19 infection. Elife 10, e63502. https://doi.org/10.7554/eLife.63502.

Müller, M., Gfeller, D., Coukos, G., and Bassani-Sternberg, M. (2017). “Hotspots” of Antigen Presentation Revealed by Human Leukocyte Antigen Ligandomics for Neoantigen Prioritization. Frontiers in Immunology 8, 1367..

Nolan, S., Vignali, M., Klinger, M., Dines, J.N., Kaplan, I.M., Svejnoha, E., Craft, T., Boland, K., Pesesky, M., Gittelman, R.M., et al. (2020). A large-scale database of T-cell receptor beta (TCRβ) sequences and binding associations from natural and synthetic exposure to SARS-CoV-2. Res Sq rs.3.rs-51964. https://doi.org/10.21203/rs.3.rs-51964/v1.

O’Donnell, T.J., Rubinsteyn, A., and Laserson, U. (2020). MHCflurry 2.0: Improved Pan-Allele Prediction of MHC Class I-Presented Peptides by Incorporating Antigen Processing. Cell Syst 11, 42–48.e7. https://doi.org/10.1016/j.cels.2020.06.010.

Ott, P.A., Hu, Z., Keskin, D.B., Shukla, S.A., Sun, J., Bozym, D.J., Zhang, W., Luoma, A., Giobbie-Hurder, A., Peter, L., et al. (2017). An immunogenic personal neoantigen vaccine for patients with melanoma. Nature 547, 217–221..

Parkhurst, M., Gros, A., Pasetto, A., Prickett, T., Crystal, J.S., Robbins, P., and Rosenberg, S.A. (2017). Isolation of T-Cell Receptors Specifically Reactive with Mutated Tumor-Associated Antigens from Tumor-Infiltrating Lymphocytes Based on CD137 Expression. Clin Cancer Res 23, 2491–2505. https://doi.org/10.1158/1078-0432.CCR-16-2680.

Pataskar, A., Champagne, J., Nagel, R., Kenski, J., Laos, M., Michaux, J., Pak, H.S., Bleijerveld, O.B., Mordente, K., Navarro, J.M., et al. (2022). Tryptophan depletion results in tryptophan-to-phenylalanine substitutants. Nature https://doi.org/10.1038/s41586-022-04499-2.

Peters, B., Nielsen, M., and Sette, A. (2020). T Cell Epitope Predictions. Annu Rev Immunol 38, 123–145. https://doi.org/10.1146/annurev-immunol-082119-124838.

Pyke, R.M., Mellacheruvu, D., Dea, S., Abbott, C.W., Zhang, S.V., Phillips, N.A., Harris, J., Bartha, G., Desai, S., McClory, R., et al. (2021). Precision Neoantigen Discovery Using Large-scale Immunopeptidomes and Composite Modeling of MHC Peptide Presentation. Mol Cell Proteomics 20, 100111. https://doi.org/10.1016/j.mcpro.2021.100111.

Reynisson, B., Alvarez, B., Paul, S., Peters, B., and Nielsen, M. (2020). NetMHCpan-4.1 and NetMHCIIpan-4.0: improved predictions of MHC antigen presentation by concurrent motif deconvolution and integration of MS MHC eluted ligand data. Nucleic Acids Res. 48, W449–W454. https://doi.org/10.1093/nar/gkaa379.

Rosenberg, S.A., and Restifo, N.P. (2015). Adoptive cell transfer as personalized immunotherapy for human cancer. Science (New York, N.Y.) 348, 62–68..

Sahin, U., Derhovanessian, E., Miller, M., Kloke, B.-P., Simon, P., Löwer, M., Bukur, V., Tadmor, A.D., Luxemburger, U., Schrörs, B., et al. (2017). Personalized RNA mutanome vaccines mobilize poly-specific therapeutic immunity against cancer. Nature 547, 222–226..

Sahin, U., Oehm, P., Derhovanessian, E., Jabulowsky, R.A., Vormehr, M., Gold, M., Maurus, D., Schwarck-Kokarakis, D., Kuhn, A.N., Omokoko, T., et al. (2020). An RNA vaccine drives immunity in checkpoint-inhibitor-treated melanoma. Nature https://doi.org/10.1038/s41586-020-2537-9.

Saini, S.K., Hersby, D.S., Tamhane, T., Povlsen, H.R., Amaya Hernandez, S.P., Nielsen, M., Gang, A.O., and Hadrup, S.R. (2021). SARS-CoV-2 genome-wide T cell epitope mapping reveals immunodominance and substantial CD8+ T cell activation in COVID-19 patients. Sci Immunol 6, eabf7550. https://doi.org/10.1126/sciimmunol.abf7550.

Sarkizova, S., Klaeger, S., Le, P.M., Li, L.W., Oliveira, G., Keshishian, H., Hartigan, C.R., Zhang, W., Braun, D.A., Ligon, K.L., et al. (2019). A large peptidome dataset improves HLA class I epitope prediction across most of the human population. Nat Biotechnol https://doi.org/10.1038/s41587-019-0322-9.

Schmidt, J., Smith, A.R., Magnin, M., Racle, J., Devlin, J.R., Bobisse, S., Cesbron, J., Bonnet, V., Carmona, S.J., Huber, F., et al. (2021). Prediction of neo-epitope immunogenicity reveals TCR recognition determinants and provides insight into immunoediting. Cell Rep Med 2, 100194. https://doi.org/10.1016/j.xcrm.2021.100194.

Shao, X.M., Bhattacharya, R., Huang, J., Sivakumar, I.K.A., Tokheim, C., Zheng, L., Hirsch, D., Kaminow, B., Omdahl, A., Bonsack, M., et al. (2020). High-Throughput Prediction of MHC Class I and II Neoantigens with MHCnuggets. Cancer Immunol Res 8, 396–408. https://doi.org/10.1158/2326-6066.CIR-19-0464.

Shimizu, K., Iyoda, T., Sanpei, A., Nakazato, H., Okada, M., Ueda, S., Kato-Murayama, M., Murayama, K., Shirouzu, M., Harada, N., et al. (2021). Identification of TCR repertoires in functionally competent cytotoxic T cells cross-reactive to SARS-CoV-2. Commun Biol 4, 1365. https://doi.org/10.1038/s42003-021-02885-6.

Shugay, M., Britanova, O.V., Merzlyak, E.M., Turchaninova, M.A., Mamedov, I.Z., Tuganbaev, T.R., Bolotin, D.A., Staroverov, D.B., Putintseva, E.V., Plevova, K., et al. (2014). Towards error-free profiling of immune repertoires. Nat Methods 11, 653–655. https://doi.org/10.1038/nmeth.2960.

Stranzl, T., Larsen, M.V., Lundegaard, C., and Nielsen, M. (2010). NetCTLpan: panspecific MHC class I pathway epitope predictions. Immunogenetics 62, 357–368. https://doi.org/10.1007/s00251-010-0441-4.

Tarke, A., Sidney, J., Kidd, C.K., Dan, J.M., Ramirez, S.I., Yu, E.D., Mateus, J., da Silva Antunes, R., Moore, E., Rubiro, P., et al. (2021). Comprehensive analysis of T cell immunodominance and immunoprevalence of SARS-CoV-2 epitopes in COVID-19 cases. Cell Rep Med 2, 100204. https://doi.org/10.1016/j.xcrm.2021.100204.

Tran, E., Ahmadzadeh, M., Lu, Y.-C., Gros, A., Turcotte, S., Robbins, P.F., Gartner, J.J., Zheng, Z., Li, Y.F., Ray, S., et al. (2015). Immunogenicity of somatic mutations in human gastrointestinal cancers. Science 350, 1387–1390. https://doi.org/10.1126/science.aad1253.

Trolle, T., McMurtrey, C.P., Sidney, J., Bardet, W., Osborn, S.C., Kaever, T., Sette, A., Hildebrand, W.H., Nielsen, M., and Peters, B. (2016). The Length Distribution of Class I-Restricted T Cell Epitopes Is Determined by Both Peptide Supply and MHC Allele-Specific Binding Preference. J. Immunol. 196, 1480–1487. https://doi.org/10.4049/jimmunol.1501721.

Viganò, S., Utzschneider, D.T., Perreau, M., Pantaleo, G., Zehn, D., and Harari, A. (2012). Functional Avidity: A Measure to Predict the Efficacy of Effector T Cells? Clinical and Developmental Immunology 2012, 1–14. https://doi.org/10.1155/2012/153863.

Vita, R., Mahajan, S., Overton, J.A., Dhanda, S.K., Martini, S., Cantrell, J.R., Wheeler, D.K., Sette, A., and Peters, B. (2019). The Immune Epitope Database (IEDB): 2018 update. Nucleic Acids Res. 47, D339–D343. https://doi.org/10.1093/nar/gky1006.

Wagih, O. (2017). ggseqlogo: a versatile R package for drawing sequence logos. Bioinformatics 33, 3645–3647. https://doi.org/10.1093/bioinformatics/btx469.

Wells, D.K., van Buuren, M.M., Dang, K.K., Hubbard-Lucey, V.M., Sheehan, K.C.F., Campbell, K.M., Lamb, A., Ward, J.P., Sidney, J., Blazquez, A.B., et al. (2020). Key Parameters of Tumor Epitope Immunogenicity Revealed Through a Consortium Approach Improve Neoantigen Prediction. Cell 183, 818–834.e13. https://doi.org/10.1016/j.cell.2020.09.015.

