## Supplementary Material for "Predictions of immunogenicity reveal potent SARS-CoV-2 CD8+ T-cell epitopes"

### Supplementary Figures

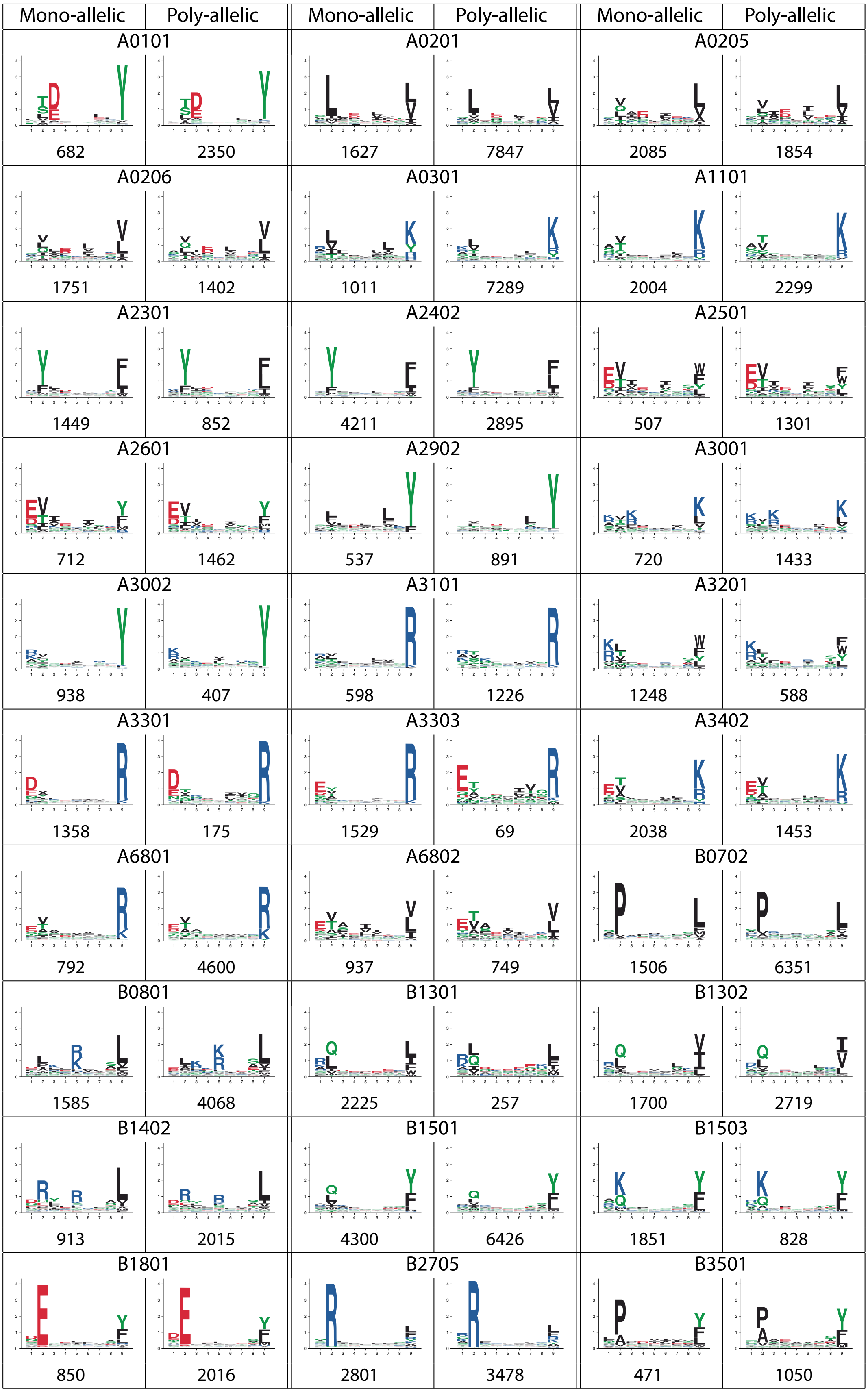

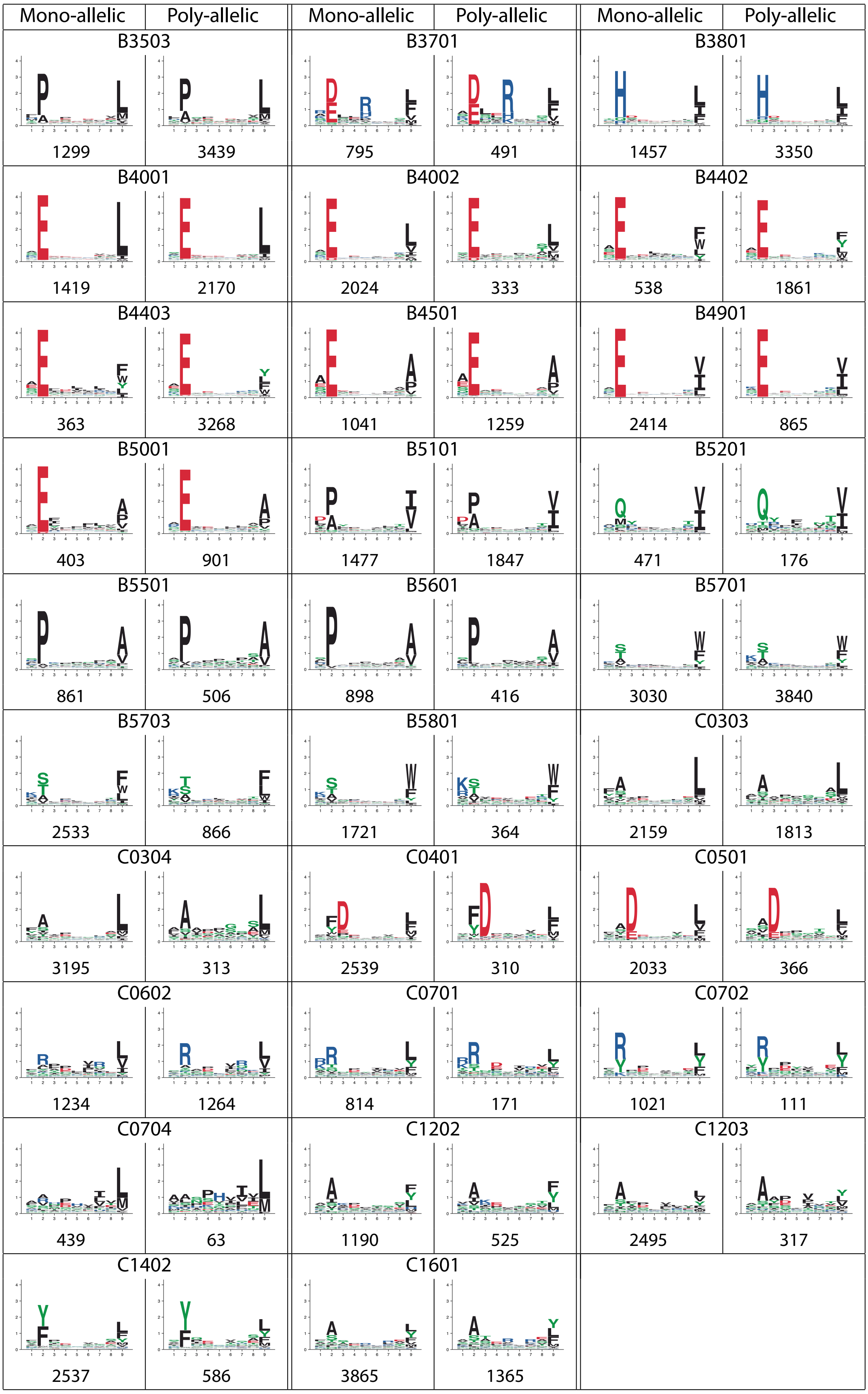


**Figure S1: Comparison between motifs deconvolved from mono-allelic and poly-allelic HLA-I peptidomics data.** All alleles found in both mono-allelic and poly-allelic samples are shown (59 in total). Numbers below each motif indicate the number of peptides (9-mers).

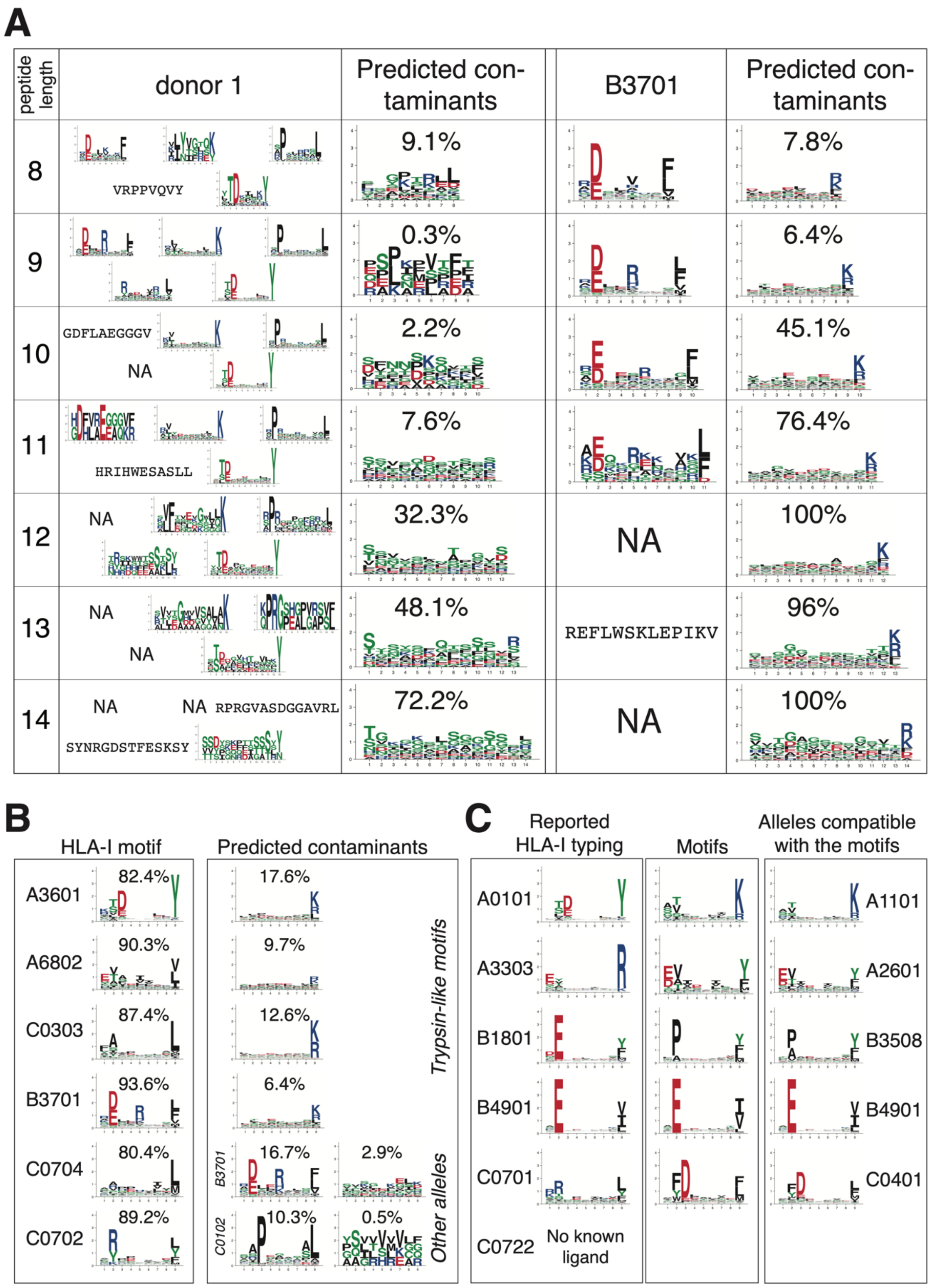


**Figure S2: Motif deconvolution predicts contaminants in mono-allelic and poly-allelic HLA-I peptidomics samples.** (A) Examples of contaminants predicted by motif deconvolution with MixMHCp for different peptide lengths in the poly-allelic sample 'donor1' (Ritz et al., 2017) and the mono-allelic sample 'B3701' (Sarkizova et al., 2019). When only one peptide is found for a given length, the sequence of the peptide is written instead of the motif. Cases without any peptide are indicated with ‘NA’. (B) Examples of different types of contaminants predicted by motif deconvolution with MixMHCp in mono-allelic samples for 9-mer peptides. Predicted contaminants include both peptides with trypsin-like protease motifs (top four) and peptides from other alleles (bottom two, contaminating alleles are indicated in italic). Data for A6802 come from Abelin et al. (Abelin et al., 2017). Data for other alleles come from Sarkizova et al. (Sarkizova et al., 2019). (C) Example of putatively erroneous HLA-I typing (sample ‘1180157F’ in (Pyke et al., 2021)). The left box shows the motifs of the alleles reported in the original study. The middle box shows the motifs identified by motif deconvolution with MixMHCp. The right box shows the motifs of alleles compatible with those predicted by motif deconvolution.

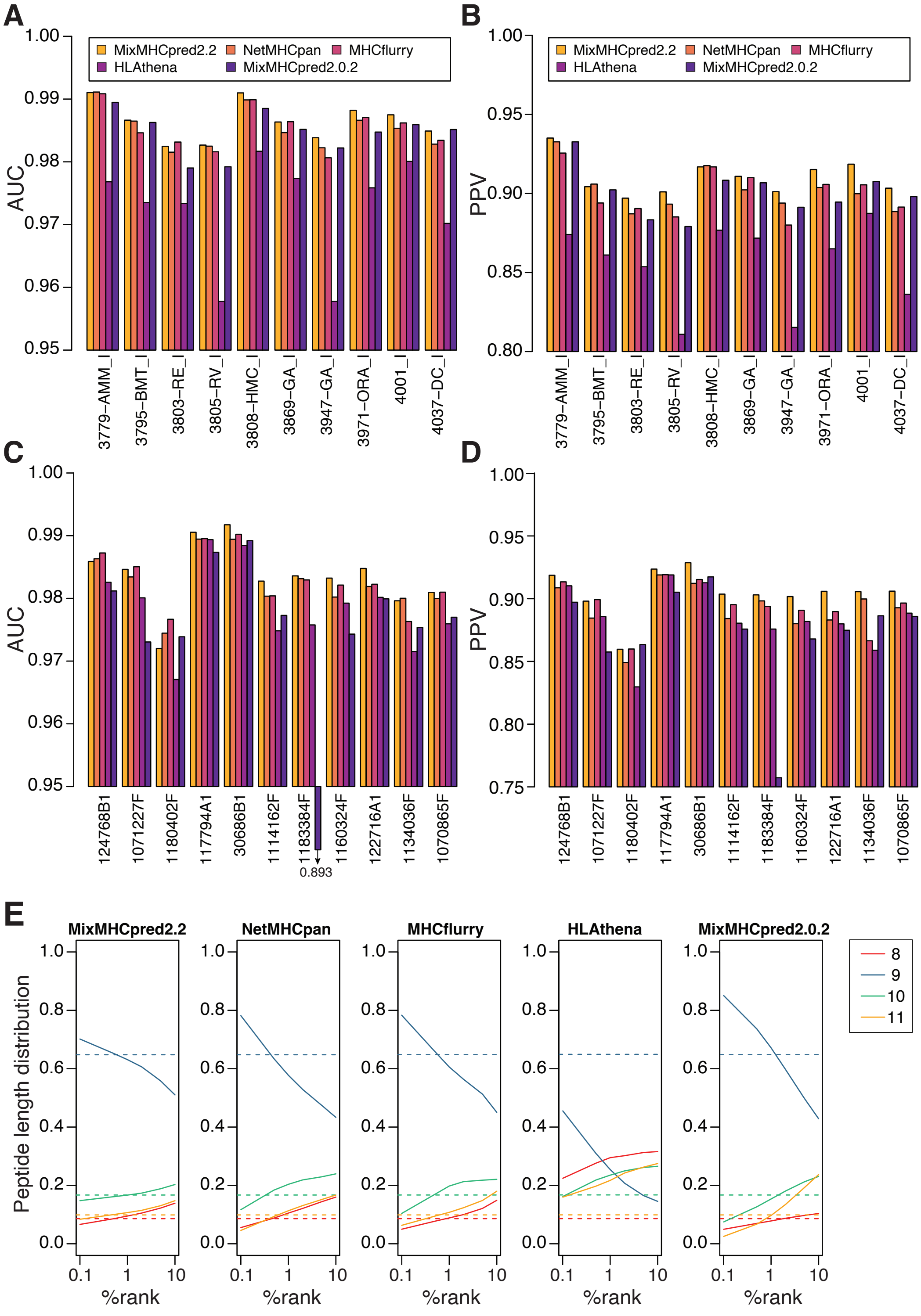


**Figure S3: Models of HLA-I binding specificities and peptide length distributions improve predictions of naturally presented HLA-I ligands.** (A-B) AUC (A) and PPV (B) values for different predictors applied on 10 HLA-I peptidomics samples from (Gfeller et al., 2018). (C-D) AUC (C) and PPV (D) values for different predictors applied on 11 HLA-I peptidomics samples from (Pyke et al., 2021). (E) Predicted peptide length distributions at multiple %rank thresholds for each HLA-I ligand predictor (average over all alleles with mono-allelic HLA-I peptidomics data). Dashed lines show the peptide length distributions observed in naturally presented HLA-I ligands from mono-allelic samples.

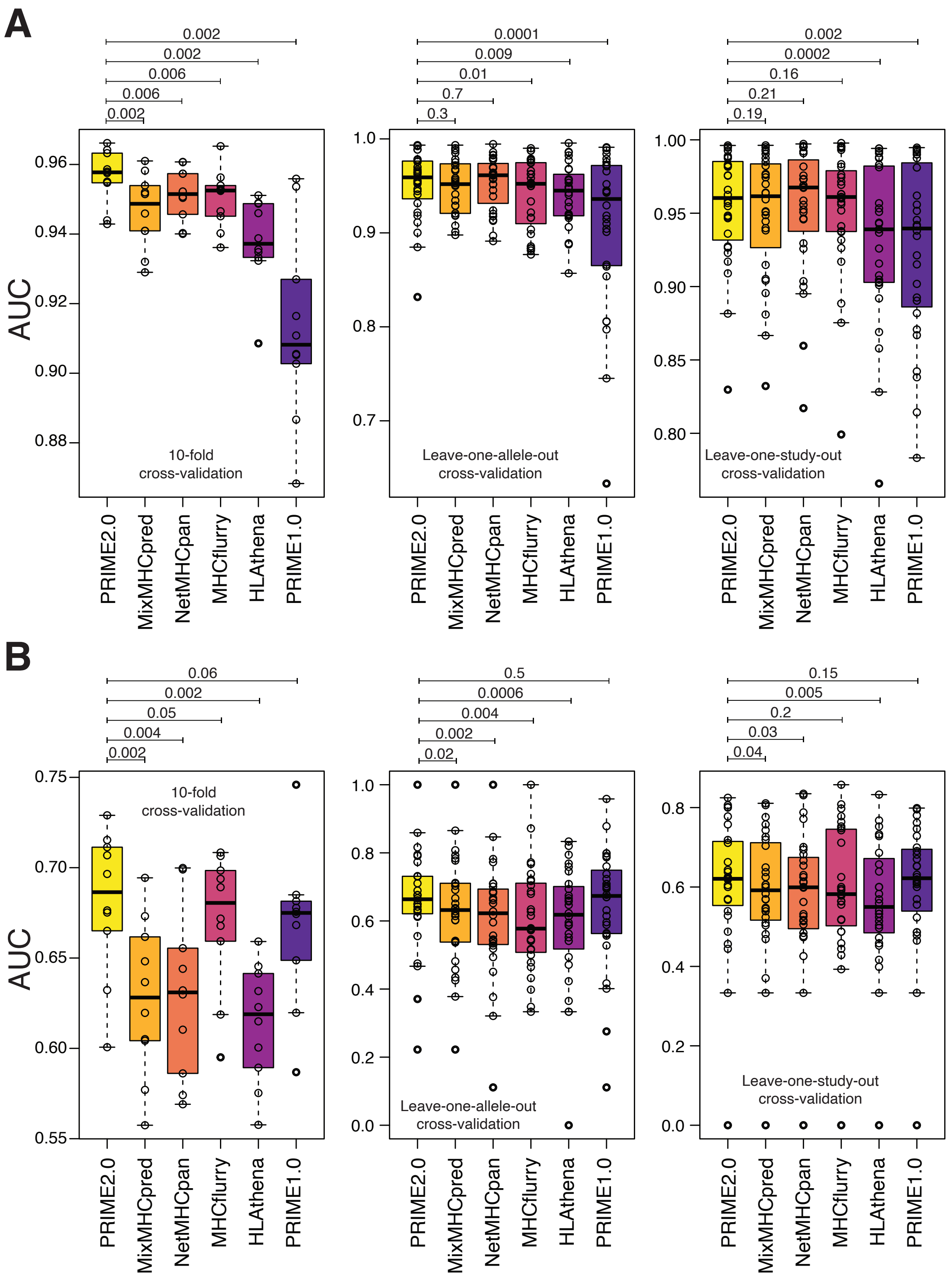


**Figure S4: Models of TCR recognition improve predictions of neo-epitopes.** (A) Details of the benchmarking of PRIME2.0 based on 10-fold cross-validation, leave-one-allele-out cross-validation and leave-one-study-out cross-validation. Boxplots show the median and lower/upper quartiles. P-values are computed with paired Wilcoxon test. (B) Same validation as in (A) after excluding randomly generated negatives in the test sets.

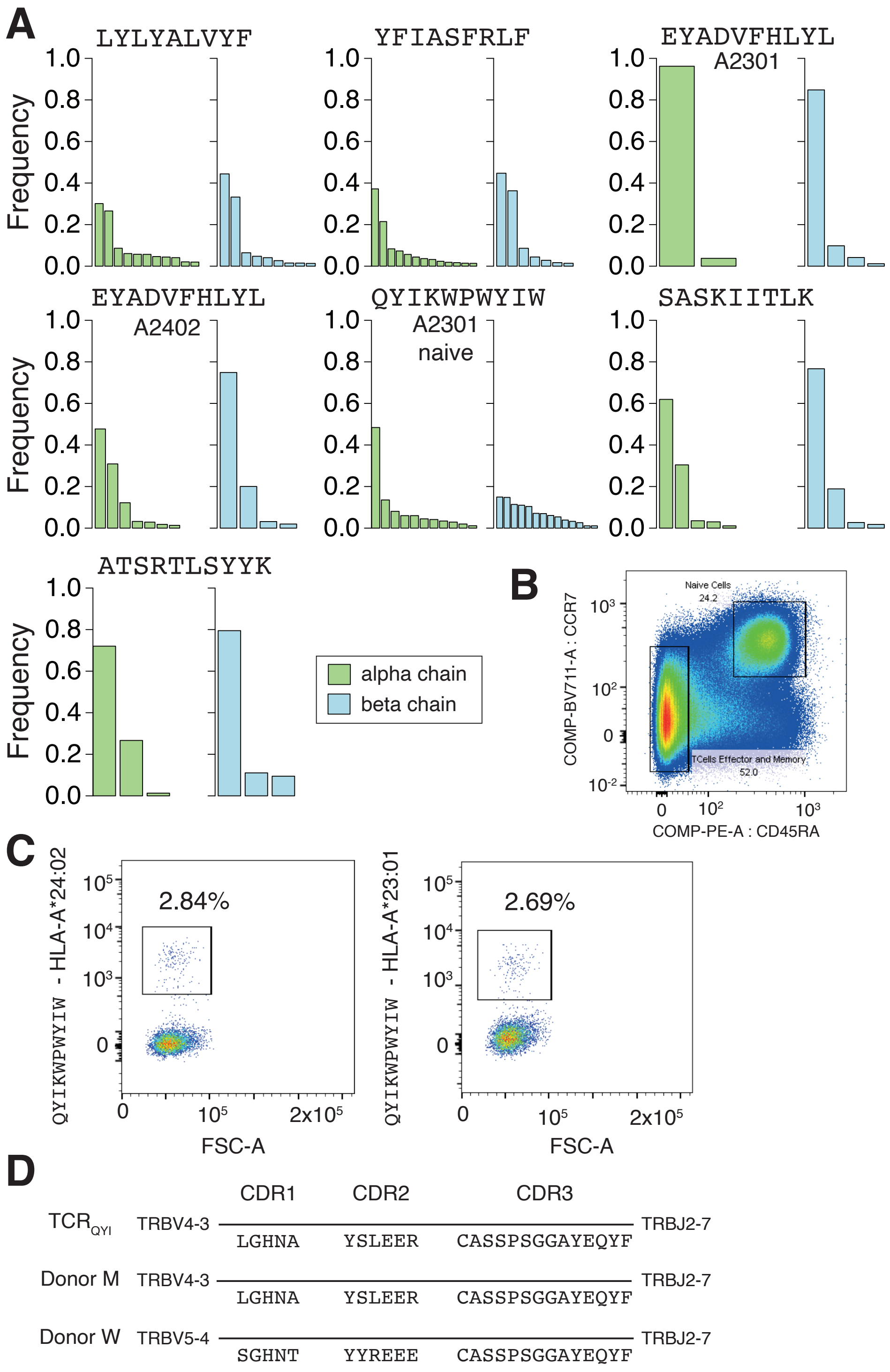


**Figure S5: Immunogenicity predictions reveal SARS-CoV-2 CD8$^+$ T-cell epitopes.** (A) Frequency of alpha and beta chains found for each epitope with more than one alpha and beta chain. (B) Sorting of naïve and effector/memory CD8$^+$ T cells in Leu184. (C) Staining of effector/memory CD8$^+$ T cells from Leu184 donor with the QYIKWPWYIW – HLA-A*24:02 and QYIKWPWYIW – HLA-A*23:01 multimers. (D) Comparison between the sequence of the TCRβ chain found in the monoclonal CD8$^+$ T cell population specific for the QYIKWPWYIW epitope (TCR$_{QYI}$) and the closest TCRβ chains found in the two COVID-

$19^+$ patients analyzed in Minervina et al. (Minervina et al., 2021) (both donors had TCRα chains identical to the one in $TCR_{QYI}$).

#### Supplementary Datasets

**Dataset S1:** List of the 244 HLA-I peptidomics samples considered in this work.

**Dataset S2:** List of HLA-I ligands used to train MixMHCpred2.2.

**Dataset S3:** List of immunogenic and non-immunogenic peptides used to train PRIME2.0

**Dataset S4: (**A) List of 213 peptides from the SARS-CoV-2 proteome included in the peptide pool. (B) List of the 15 most common HLA-I alleles used to make predictions. (C) Information about the six donors used to screen SARS-CoV-2 peptides for immunogenicity.

**Dataset S5:** List of TCR sequences and UMI counts (alpha and beta chains) obtained from $CD8^+$ T cells recognizing seven SARS-CoV-2 epitopes.

### References

Abelin, J.G., Keskin, D.B., Sarkizova, S., Hartigan, C.R., Zhang, W., Sidney, J., Stevens, J., Lane, W., Zhang, G.L., Eisenhaure, T.M., et al. (2017). Mass Spectrometry Profiling of HLA-Associated Peptidomes in Mono-allelic Cells Enables More Accurate Epitope Prediction. Immunity *46*, 315–326. https://doi.org/10.1016/j.immuni.2017.02.007.

Gfeller, D., Guillaume, P., Michaux, J., Pak, H.-S., Daniel, R.T., Racle, J., Coukos, G., and Bassani-Sternberg, M. (2018). The Length Distribution and Multiple Specificity of Naturally Presented HLA-I Ligands. J. Immunol. *201*, 3705–3716. https://doi.org/10.4049/jimmunol.1800914.

Minervina, A.A., Komech, E.A., Titov, A., Bensouda Koraichi, M., Rosati, E., Mamedov, I.Z., Franke, A., Efimov, G.A., Chudakov, D.M., Mora, T., et al. (2021). Longitudinal high-throughput TCR repertoire profiling reveals the dynamics of T-cell memory formation after mild COVID-19 infection. Elife *10*, e63502. https://doi.org/10.7554/eLife.63502.

Pyke, R.M., Mellacheruvu, D., Dea, S., Abbott, C.W., Zhang, S.V., Phillips, N.A., Harris, J., Bartha, G., Desai, S., McClory, R., et al. (2021). Precision Neoantigen Discovery Using Large-scale Immunopeptidomes and Composite Modeling of MHC Peptide Presentation. Mol Cell Proteomics *20*, 100111. https://doi.org/10.1016/j.mcpro.2021.100111.

Ritz, D., Gloger, A., Neri, D., and Fugmann, T. (2017). Purification of soluble HLA class I complexes from human serum or plasma deliver high quality immuno peptidomes required for biomarker discovery. Proteomics *17*. https://doi.org/10.1002/pmic.201600364.

Sarkizova, S., Klaeger, S., Le, P.M., Li, L.W., Oliveira, G., Keshishian, H., Hartigan, C.R., Zhang, W., Braun, D.A., Ligon, K.L., et al. (2019). A large peptidome dataset improves HLA class I epitope prediction across most of the human population. Nat Biotechnol https://doi.org/10.1038/s41587-019-0322-9.
